# PHYTOCHROME INTERACTING FACTOR 7 moderates the activity of phytochrome A in canopy shade conditions

**DOI:** 10.1101/2025.10.07.680213

**Authors:** Mengke Zhou, Andrés Romanowski, Karen J. Halliday

**Author notes:** Corresponding authors: Mengke Zhou and Karen J. Halliday, Institute for Molecular Plant Sciences, School of Biological Sciences, University of Edinburgh, Edinburgh EH9 3BF, United Kingdom.

## Abstract

In nature, plants perceive and respond to complex vegetative environments through the phytochrome (phy) photoreceptor-signaling system. The canopy adaptation strategy, elicited by phyA, together with the shade avoidance syndrome, regulated by phyB, enables plants to adapt to diverse forms of vegetation shade. The phyA High Irradiance Response (HIR) mode, which involves photocycle-coupled nuclear shuttling, is essential for sensing sustained far-red rich conditions typical of canopy or occlusive shade. Our research identifies PHYTOCHROME INTERACTION FACTOR 7 (PIF7) as a key regulator of phyA function. We show that PIF7ox effectively suppresses phyA action. Under persistent shade, PIF7 and phyA both accumulate and interact within nuclear photobodies, with binding mediated by the C-terminal domain of PIF7 and the PAS-PAS domain of phyA. PIF7 also interacts with FHY1 and FHL transport facilitators, thereby interfering with phyA cytosolic-nuclear shuttling. This coordinated molecular targeting by PIF7 constitutes an effective strategy for modulating phyA function under shaded conditions.

## INTRODUCTION

Light sensing is fundamental for plant adaptation and survival in vegetative shade. This is particularly important during seedling establishment, a stage when plants are most vulnerable. Plants have evolved distinct developmental strategies to cope with different types of shade that are prevalent in nature. The light receptor phytochrome B (phyB), is able to detect FR-rich light from neighboring plants and elicit the Shade Avoidance Syndrome (SAS), which induces elongation of seedling hypocotyls, repositions leaves to improve light capture, and triggers early flowering (*1,2*). Under persistent canopy shade, plants adopt alternative strategies, such as the phyA-mediated Canopy Adaptation Strategy (CAS), which suppresses hypocotyl elongation and conserves resources, optimizing growth when shade is unyielding (*3*). As vegetative habitats can change considerably through a season, the ability to adjust photoreceptor activity offers adaptive flexibility, ensuring successful seedling establishment and progression through the life cycle. In Arabidopsis, phytochromes A-E, form a family of photoreceptors that principally absorb light in the red (R) and far-red (FR) regions of the spectrum (*4*). Absorption of R photoconverts phytochrome into its active Pfr form, while FR photoconverts Pfr to the inactive Pr conformer. When activated phytochromes move from the cytosol to the nucleus where they aggregate in sub-nuclear photobodies to perform a range of functions (*5,6*). While all phytochromes possess similar photosensory properties, they fall into two classes based on their distinct modes of action. Phytochromes B-E, which operate in the low fluence response (LFR) mode, are activated by R and deactivated by FR light. Consequently, in FR-rich vegetation shade environments, these phytochromes are less active (*4,7*). A key distinction of phyA is that it is activated by FR-rich light through its high irradiance response (HIR) mode of action (*7*). This property arises partly from its photobiology and partly from the shuttling mechanism that concentrates phyA in the nucleus (*8*). PhyA lacks a nuclear localization signal and primarily relies on the proteins FHY1/FHL to facilitate its transport into the nucleus. These proteins specifically recognize phyA-Pfr and transport it to the nucleus, where it is converted into phyA-Pr, allowing FHY1/FHL to dissociate and be recycled to the cytoplasm for further shuttling (*8-10*). Nuclear located phyA regulates a number of processes, including expression of target genes, through association with transcription factors such as PHYTOCHROME INTERACTING FACTORS (PIFS) and ELONGATED HYPOCOTYL 5 (HY5) (*11*). The central role of FHY1/FHL shuttling in the phyA HIR, identifies this mechanism as a potential target for modifying phyA action and the adaptive response (*8*). Indeed, HY5 has been reported to be involved in negative-feedback regulation of phyA signaling through repressing FHY1/FHL expression (*11*). Here, HY5 directly interacts with the DNA-binding regions of FAR-RED ELONGATED HYPOCOTYLS 3 (FHY3) and FAR-RED IMPAIRED RESPONSE 1 (FAR1) to prevent their transcriptional activation of FHY1/FHL.

PIF transcription factors are well-studied negative regulators of phytochrome signaling. They function by directly binding G/PBE-boxes in the promoters of their target genes (*12*). In Arabidopsis, there are eight PIFs (PIF1, PIF2/PIL1-PIF8), each with an active phytochrome B (APB) domain that facilitates their interaction with phyB. Only PIF1 and PIF3 contain an active phytochrome A (APA) domain, which directly binds the C-terminus of phyA, comprising two Period-Arnt-SIM (PAS) and a histidine kinase-related (HKRD) domain (*13*). In FR light, phyA directly interacts with PIF1 and PIF3, triggering their proteolytic destruction and/or sequestration from target promoters (*13,14*). Although PIF4, PIF5, PIF8 and PIF7 lack an APA domain, they have been shown to control aspects of phyA signaling (*15-17*). Interestingly, levels of PIF8 protein are boosted by phyA. In darkness, CONSTITUTIVE PHOTOMORPHOGENESIS 1 (COP1) promotes PIF8 protein degradation through the 26S proteasome, thereby reducing its abundance. Conversely, under FR light, phyA inhibition of COP1 stabilizes the PIF8 protein, forming a negative feedback loop that limits phyA action (*15*).

Notably, PIF7, a close homologue of PIF8, is a potent regulator of SAS. In unshaded conditions, PIF7 interacts with active phyB (Pfr) through its APB domain, and both proteins localize to sub-nuclear photobodies. This interaction prevents PIF7 from binding to the promoters of target genes, effectively inhibiting PIF7-regulated gene expression (*18,19*). Under shade conditions, the deactivation of phyB releases PIF7 from this repression, leading to the activation of shade-responsive genes and triggering the physiological SAS response (*20,21*). Furthermore, exposure to shade triggers the dephosphorylation and nuclear accumulation of PIF7, processes that enhance the activation of SAS (*20*). Given its strong promotory role in SAS, it is notable that PIF7 has recently been demonstrated to increase *PHYA* transcript abundance, an action that could potentially limit SAS. This PIF7 action can be inhibited by *PHYA UTR ANTISENSE RNA (PUAR)*, which binds to PIF7 and prevents its interaction with *PHYA’*s 5’ UTR, thereby reducing *PHYA* transcript levels (*17*). Collectively, this research underscores the importance of PIF7 as a key agent in executing and moderating the molecular response to shade.

Here, we show that PIF7 modulates phyA signaling under vegetative shade by controlling phyA subcellular localization and activity. PIF7 overexpression (PIF7ox) abolishes phyA-mediated suppression of seedling hypocotyl elongation under persistent shade. Under these conditions, PIF7 and phyA co-localize in nuclear photobodies and interact directly via the PIF7 prion-like domain and the phyA C-terminus. Additionally, our data shows that PIF7 governs phyA subcellular localization by modulating FHY1/FHL-dependent nuclear import. These insights clarify how PIF7 effectively restricts both the nuclear transport and signaling activities of phyA, illuminating the intricate molecular mechanisms at play.

## RESULTS

### PIF7ox blocks phyA action

PIF7, which functions downstream of phyB, also shares significant sequence similarity with PIF8, a regulator of phyA responses. This similarity led us to investigate the potential genetic interaction between PIF7 and phyA. To achieve this, we measured the hypocotyl length of 6-day-old seedlings grown under two regimes: 8-hour light (PAR = 85 ± 5 μmol m^−2^ s^−1^): 16-hour dark cycle (WL) at 22°C, or identical conditions with continuous far-red (FR) light supplied through the photoperiod (WLFR) (Fig. 1A). This second regime was designed to simulate persistent moderate shade, delivering a red:far-red (R:FR) ratio of 0.2 and Pfr/Ptot of 0.27 (fig. S1 and table S1). In all experiments, supplementary FR light was applied from one day after germination to align with the optimal period for phytochrome A (phyA) action during seedling development (*3,22*). As expected, exposure to WLFR inhibited hypocotyl elongation in wild-type (WT, Col-0) seedlings, whereas *phyA-211* mutants instead showed enhanced elongation in WLFR (Fig. 1B). This pronounced elongation in *phyA-211* reflects the loss of phyA-mediated repression of SAS, which is predominantly initiated by phyB inactivation under WLFR conditions (*22,23*) The phyAox line exhibited a constitutively short hypocotyl, which was further reduced under WLFR (Fig. 1B). Together, these results confirm that our simulated shade growth regime effectively activates phyA-dependent inhibition of hypocotyl elongation and robustly counteracts SAS-associated growth.

**Fig. 1.**
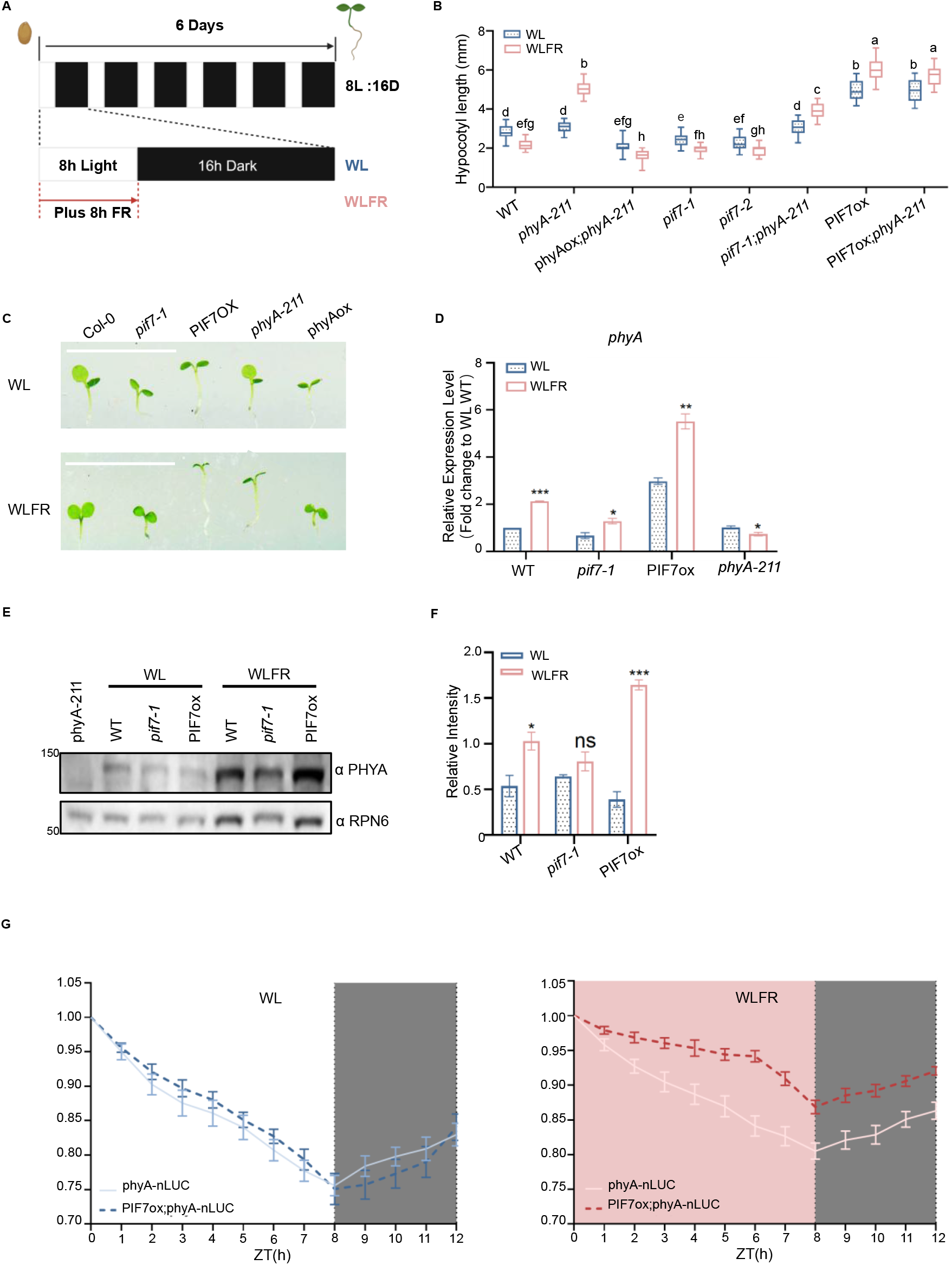
PIF7ox moderates phyA function under WLFR. (**A**) Schematic of experimental growth conditions. Seeds were sown on 1/2 MS plates and exposed to 4 hours (h) of light to induce germination, followed by 20 h of dark. Seedlings were then grown for 6 days in white light (WL) photoperiods with or without supplementary far-red (WLFR), lowering the R:FR to 0.20 (red arrow). All experiments were conducted at 22 °C under a photosynthetically active radiation (PAR) fluence rate of 85 ± 5 μmol m^−2^ s^−1^. (**B**) Hypocotyl length (mm) of WT (Col-0), *phyA-211, p35S::PHYA-tagRFP:terRbcS;phyA-211* (phyAox), *pif7-1, pif7-2, pif7-1;phyA-211, p35S::PIF7ox-FLASH* (PIF7ox) and PIF7ox*;phyA-211*. Seedlings were grown for 6 days in WL or WLFR. (**C**) Representative images of WT (Col-0), *pif7-1*, PIF7ox, phyAox and *phyA-211* seedlings grown in WL or WLFR for 6 days. Scale bar = 1 cm. (**D**) Real-time qPCR was used to quantify *PHYA* mRNA levels in WT (Col-0), *pif7-1*, PIF7ox and *phyA-211* seedlings using *PP2A* as the internal control. Samples were collected at ZT7 (1 hour before dusk) from 6d-old seedlings grown in WL or WLFR. (**E** and **F**) PHYA protein levels in WT (Col-0), *pif7-1* and PIF7ox were determined by immunoblotting. Anti-PHYA was used to detect native phyA, while anti-RPN6 served as an internal control. The bar chart (**F**) displays the relative levels of phyA, normalised to RPN6, based on data from four biological replicates collected at ZT7. (**G**) Diel phyA-nLUC levels were quantified in WT (Col-0), *pif7-1* and PIF7ox lines. The phyA-nLUC line was generated by expressing (p*PHYA::PHYA-nLUC*) in *phyA-211*, while the PIF7ox;phyA-nLUC was generated by introgressing *pPHYA::PHYA-nLUC* into the PIF7ox line. Bioluminescence signals were recorded from day 6. For (**B, D, F** and **G**) data are presented as mean values ± s.e.m., for *n*=25 (seedling numbers) in (B); and for *n*=3,4 and 9 biological repeats in (**D, F** and **G**), respectively. For (**B**) a Two-way ANOVA was conducted to assess statistical significance across light conditions for each genotype (α=0.05), followed by Tukey’s HSD post hoc test for pairwise multiple comparisons. Groups that do not differ significantly are indicated by the same letter. For (**D** and **F**) Asterisks indicate significant differences from the WL values (*, P < 0.05; **, P < 0.01; and ***, P < 0.001; Student’s t test).

We also found that similar to phyAox, the loss-of-function alleles *pif7-1* and *pif7-2* exhibited slightly shorter hypocotyls than the WT while maintaining responsiveness to WLFR (Fig. 1, B and C). These phenotypic effects were abolished by the *phyA-211* mutation, as observed in the *pif7-1;phyA-211* double mutant. Consistent with the established role of PIF7 in promoting the phyB-mediated SAS, in WLFR *pif7-1* reduced hypocotyl elongation in the *phyA-211* background (*24,25*). In comparison, hypocotyls of *p35S::PIF7-FLASH* (PIF7ox) seedlings, which have 3-fold (WL) change than WT higher expression (fig. S2), were significantly elongated under WL, with WLFR inducing a further increase in elongation (Fig. 1, B and C). While PIF7ox and PIF7ox;*phyA-211* exhibited nearly identical hypocotyl responses. Our findings, therefore, provide genetic evidence linking PIF7 and phyA. This relationship is particularly evident in PIF7ox seedlings, which display exaggerated SAS-associated elongation and a complete loss of the WLFR-induced phyA response (Fig. 1, B and C). The data collectively suggests a mechanism in which PIF7 participates in a cross-regulatory interaction between phyB and phyA.

Given the capacity of PIF7ox to inhibit phyA activity, we next examined the possibility of functional redundancy with other PIFs. As anticipated, PIF8ox also lacked WLFR-mediated growth inhibition, but PIF3ox, PIF4ox and PIF5ox all retained this response (fig. S3, A and B) (*15*). Our data therefore illustrate that even though PIF3, PIF4 and PIF5 have been implicated in *PHYA* regulation and signaling, their roles are functionally distinct from those of PIF7 and PIF8 (*16,26,27*). We found the hypocotyl lengths of *pif8-1* to be comparable to WT, while *pif8-1* slightly reduced the impact of *phyA-211* on hypocotyl elongation in WLFR, as observed in *pif8-1*;*phyA-211* (fig. S3C). Further, PIF8ox;*phyA-211* hypocotyls were more elongated than PIF8ox, indicating that while PIF8ox suppresses the phyA-dependent response, it does not completely eliminate it. These findings are consistent with the reported role of PIF8 as a negative regulator of phyA signaling (*15*). However, our data also showed the genetic interaction with *phyA-211* differs between PIF8ox and PIF7ox. Whereas *phyA-211* is epistatic to PIF8ox, the relationship is reversed for PIF7ox, which is epistatic to *phyA-211* (Fig. 1, B and fig. S3C). The difference in epistasis for PIF7 likely reflects its dual role in phyA- and phyB-mediated signaling pathways. However, with respect to phyA signaling, we wanted to determine whether PIF7 has an analogous role to PIF8, or if it operates through a different mechanism.

### PIF7 moderates phyA protein abundance

Previous studies have demonstrated that *PHYA* expression is strongly induced by shade and that PIF7 increases *PHYA* transcript abundance through binding to the *PHYA* 5′ UTR (*3*,1*7*). Consistent with these findings, we observed that under an 8L:16D photoperiod, sampling one hour before dusk (ZT7), WLFR enhanced *PHYA* expression in WT seedlings, whereas this response was attenuated in the *pif7-1* mutant (Fig. 1D). In comparison, PIF7ox exhibited elevated *PHYA* levels in control conditions, which were further enhanced in WLFR.

Immunoblot analysis with an antibody against native phyA revealed consistently low phyA levels under WL across all genotypes, reflecting the well-established light lability of phyA and its degradation upon white light exposure (*3,28*) ^(^Fig. 1, E and F). Indeed, we also found that PIF7ox had minimal effect on *pPHYA::PHYA-nLUC* (phyA-nLUC) activity through a WL photoperiod, where reporter levels gradually decline over the course of the day (Fig. 1G) (*3*). In contrast, under WLFR, phyA protein levels were elevated in WT, further increased in PIF7ox, and reduced in *pif7-1*, closely reflecting the corresponding patterns of *PHYA* gene expression (Fig. 1, D to F). We also observed higher levels of daytime phyA-nLUC in WLFR, which was notably elevated by PIF7ox (Fig. 1G) (*3*). Thus, our findings demonstrate that under WLFR conditions, PIF7 regulates *PHYA* expression and the abundance of phyA protein. However, despite accumulating significantly higher levels of phyA under WLFR, PIF7ox seedlings fail to elicit a phyA response. This indicates that the regulatory effect of PIF7ox on phyA activity does not stem from changes in phyA abundance.

To examine the *in vivo* dynamics of PIF7 under our experimental conditions, we generated *pPIF7::PIF7-nLUC* (PIF7-nLUC) lines. We discovered that PIF7-nLUC exhibits rhythmic patterns, with higher levels during the day compared to the night (fig. S4A). WLFR boosted daytime levels of PIF7-nLUC levels, with the *phyA-211* mutation elevating PIF7-nLUC toward the end of the period, especially in WLFR (fig. S4B). Therefore, our data indicated that similar to phyA, WLFR increases the abundance of PIF7, and that phyA plays a minor role in reducing PIF7 production toward the end of the photoperiod.

### PIF7 can interact with phyA in nuclear photobodies

To establish whether PIF7 and phyA are found in the same cellular location we analyzed transgenic lines expressing *pPHYA::PHYA*-sGFP (phyA-GFP) and *p35S:GFP-PIF7* (PIF7-GFP). Consistent with earlier studies, we found that under WL, PIF7-GFP and phyA-GFP were predominantly in the cytosol, while exposure to WLFR enhanced their nuclear localization (fig. S5)(*20,29,30*) Previously, PIF7 was shown to form sub-nuclear photobodies, or speckles, in response to red light (*18*). Similarly, we observed nuclear PIF7 photobodies in WL and discovered that PIF7 also formed photobodies under WLFR (fig. S6). As phyA-GFP locates to photobodies in WLFR, we reasoned that PIF7 and phyA may co-localize in these sub-nuclear foci (fig. S6). To investigate this, we employed the *p35S::PIF7-sfGFP* (PIF7-GFP) and *p35S::PHYA-RFP* (phyA-RFP) constructs which were transiently expressed in *N. benthamiana* leaves. After exposure to WLFR, we observed co-localization of PIF7-GFP and phyA-RFP in nuclei and in photobodies (Fig. 2A). Quantitative fluorescence measurements along a nuclear transect, intersecting two photobodies, confirmed this co-localization (Fig. 2A).

**Fig. 2.**
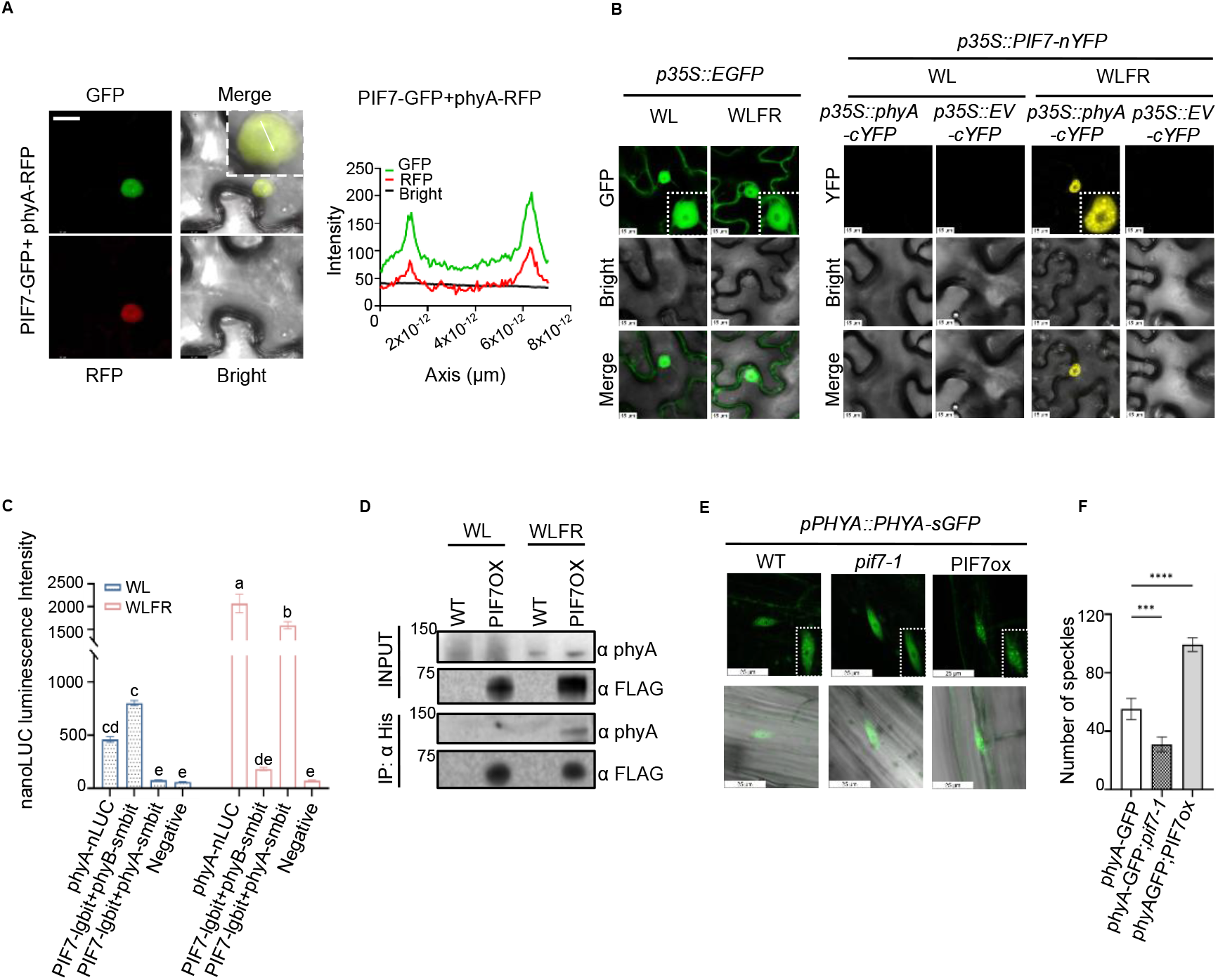
In WLFR PIF7 and phyA interact in nuclear photobodies. (**A**) Subcellular localization of *p35S::PIF7-sfGFP* (PIF7-GFP) and *p35S::PHYA-RFP* (phyA-RFP) under WLFR in *Nicotiana benthamiana* leaf epidermal cells. Co-localisation is evident from the chart (right) which traces the GFP and RFP channel signals along the white bar (intersecting two photobodies) shown in merged nucleus image (left). (**B**) Bimolecular Fluorescence Complementation (BiFC) assays confirm PIF7 and phyA interact in nuclear photobodies. BiFC was detected between *p35S::PIF7-nYFP* (PIF7-nYFP) and *p35S::PHYA*-cYFP (phyA-cYFP) in WLFR, but not with the *p35S::EV-cYFP* (EV-cYFP, empty vector) control. The *p35S::EGFP*, the transformation control, was located across cellular compartments irrespective of light conditions. All the constructs were expressed in *N. benthamiana*. Scale Bar =15μm. (**C**) Split NanoLUC (nLUC) assay demonstrates PIF7 and phyA physically interact. *p35S::*PIF7-lgbitnLUC (PIF7-lgbit), *p35S::PHYA-smbitnLUC* (phyA-smbit), *p35S::PHY*B-smbitnLUC (phyB-smbit) and *p35S::PHY*A-nLUC (phyA-nLUC) reporter constructs were expressed in *Nicotiana benthamiana*. A WLFR-dependent interaction was demonstrated for PIF7-lgbit+phyA-smbit. PhyA-nLUC was used as light conditional controls, and phyB-smbit+phyA-smbit as the negative control. (**D**) *In vivo* co-immunoprecipitation assay (Co-IP) illustrates that under WLFR, phyA interacts with PIF7 in Arabidopsis. Anti-His beads were used to pull down PIF7 from PIF7ox (*p35S::PIF7-FLASH*; 9×Myc-6×His-3×FLAG) or the WT(*Col-0*) negative control. Anti-PHYA and anti-FLAG used for detecting phyA and PIF7 respectively, in the IP/Input. Samples were collected at ZT7 from 6-d old seedlings exposed to WL or WLFR. (**E** and **F**) PIF7 involvement in PHYA photobody formation was assessed by expressing *pPHYA::PHYA-sGFP* (PHYA-GFP) in *pif7-1* and *PIF7ox* backgrounds. Representative images of hypocotyl epidermal cell nuclei are shown (E), together with quantification of nuclear photobodies from 10 nuclei per seedling across five seedlings (F). Data are presented as mean ± s.e.m. Images were captured at ZT7 after 6 days of growth under WL or WLFR. Imaging focused on the mid-hypocotyl region using a 20 × 9.91 objective lens. Scale bar = 25 μm. For **(A, B, E** and **F**), the GFP/YFP and RFP are specific channels with associated filters, signals were detected using 488nm and 552nm laser wavelengths, the Bright is bright field. For **(C**) data are presented as mean values ± s.e.m., *n*=12 (leaf numbers). A Two-way ANOVA was conducted to assess statistical significance across light conditions for each genotype (α=0.05), followed by Tukey’s HSD post hoc test for pairwise multiple comparisons. Groups that do not differ significantly are indicated by the same letter. For (**F**) A One-way ANOVA test was used to obtain statistical significance (α=0.05); the Dunnett test for multiple comparisons. Asterisks indicate significant differences from the phyA-GFP values (***, P < 0.001; and ****, P < 0.0001).

Next, we created split bimolecular fluorescence complementation (BiFC) constructs *p35S::PIF7-nYFP* (PIF7-nYFP) and *p35S::PHYA-cYFP* (phyA-cYFP), to test whether PIF7 and phyA physically interact *in vivo*. Initially, we analyzed the *p35S::EGFP* control, which was detected across cytosolic and nuclear compartments under both WL and WLFR (Fig. 2B). In contrast, our BiFC assays with PIF7-nYFP and phyA-cYFP exhibited strong signals in the nucleus and in sub-nuclear foci following exposure to WLFR. Notably, these signals were absent in control lines expressing *p35S::PIF7-nYFP* and *p35S::EV-cYFP*. The findings suggest that PIF7 and phyA can indeed interact under WLFR conditions (Fig. 2B).

To gain further evidence for this interaction we also generated split nanoLUC (nLUC) luciferase reporter constructs, *p35S::PIF7-lgbitnLUC* (PIF7-lgbit) and *p35S::PHYA-smbitnLUC* (phyA-smbit), with *p35S::PHYB-smbitnLUC* (phyB-smbit) and *p35S::PHYA-nLUC* (phyA-nLUC) as controls (Fig. 2C). As expected, the phyA-nLUC control emitted a strong signal in WLFR compared to WL, as WLFR typically enhances phyA abundance (*3*). Conversely, the PIF7-lgbit+phyB-smbit control had a strong signal in WL which diminished in WLFR, consistent with the known light conditional interaction between PIF7 and phyB (*18*). The negative controls (phyA-smbit + phyB-smbit) had low signals in both conditions. In the WL regime, the signal from PIF7-lgbit + phyA-smbit was low, similar to that of the negative control, whereas in WLFR, the signal was strong, confirming the WLFR-specific PIF7-phyA interaction. Additionally, using an antibody against native phyA, we demonstrated co-immunoprecipitation of phyA with PIF7ox, in WLFR but not WL conditions (Fig. 2D). Thus, our data provides evidence that PIF7 and phyA directly interact within photobodies after exposure to WLFR, which promotes their respective nuclear accumulation (*20,31*).

Interestingly, our data illustrates that PIF7-GFP forms photobodies in both WL and WLFR, yet phyA photobodies are only detected in our WLFR regime (fig. S6). We therefore reasoned that PIF7 may play a role in phyA photobody formation under WLFR. In support of this notion, we found that in WLFR, phyA-GFP photobodies were not detectable in *pif7-1* but were more abundant in the PIF7ox line (Fig. 2E). It therefore appears that PIF7 may be required for phyA to photobody formation in WLFR.

### The PIF7 C-terminus interacts with the phyA PAS-PAS domain

Earlier work demonstrated that phyA binds directly with PIF1 and PIF3. The phyA C-terminal domain is known to be important for this interaction, as is the active phyA-binding domains (APA) of PIF1 and PIF3(*13,26*). To identify PHYA regions that required for the molecular interaction we utilized *p35S::PIF7-nYFP* (PIF7-nYFP) and *p35S::PHYA-cYFP* (phyA-cYFP) split YFP assay, creating truncated PHYA constructs as follows: C-terminal (PHYA-C), comprising HINGE (H), PAS-PAS DOMAIN (PAS), and HISTIDINE KINASE RELATED DOMAIN (HKRD) regions [*H+PAS+HKRD-cYFP*], PHYA-C1 [*PAS+HKRD-cYFP*], PHYA-P [*PAS-cYFP*], and PHYA-H [*HKRD-cYFP*] (Fig. 3A). As before, the *p35S::EGFP* control exhibited non-specific cellular localization pattern in WL and WLFR (Fig. 2B and Fig, 3A). In contrast we obtained WLFR-specific nuclear fluorescence signals for PIF7-nYFP and the full-length phyA-cYFP control, as well as the truncated PHYA-C, PHYA-C1, and PHYA-P constructs. No signal was observed for PHYA-H and the negative EV control. The strong signal detected for PHYA-P, which includes PAS-PAS domain motifs, highlights the significance of this molecular region in facilitating the interaction with PIF7.

**Fig. 3.**
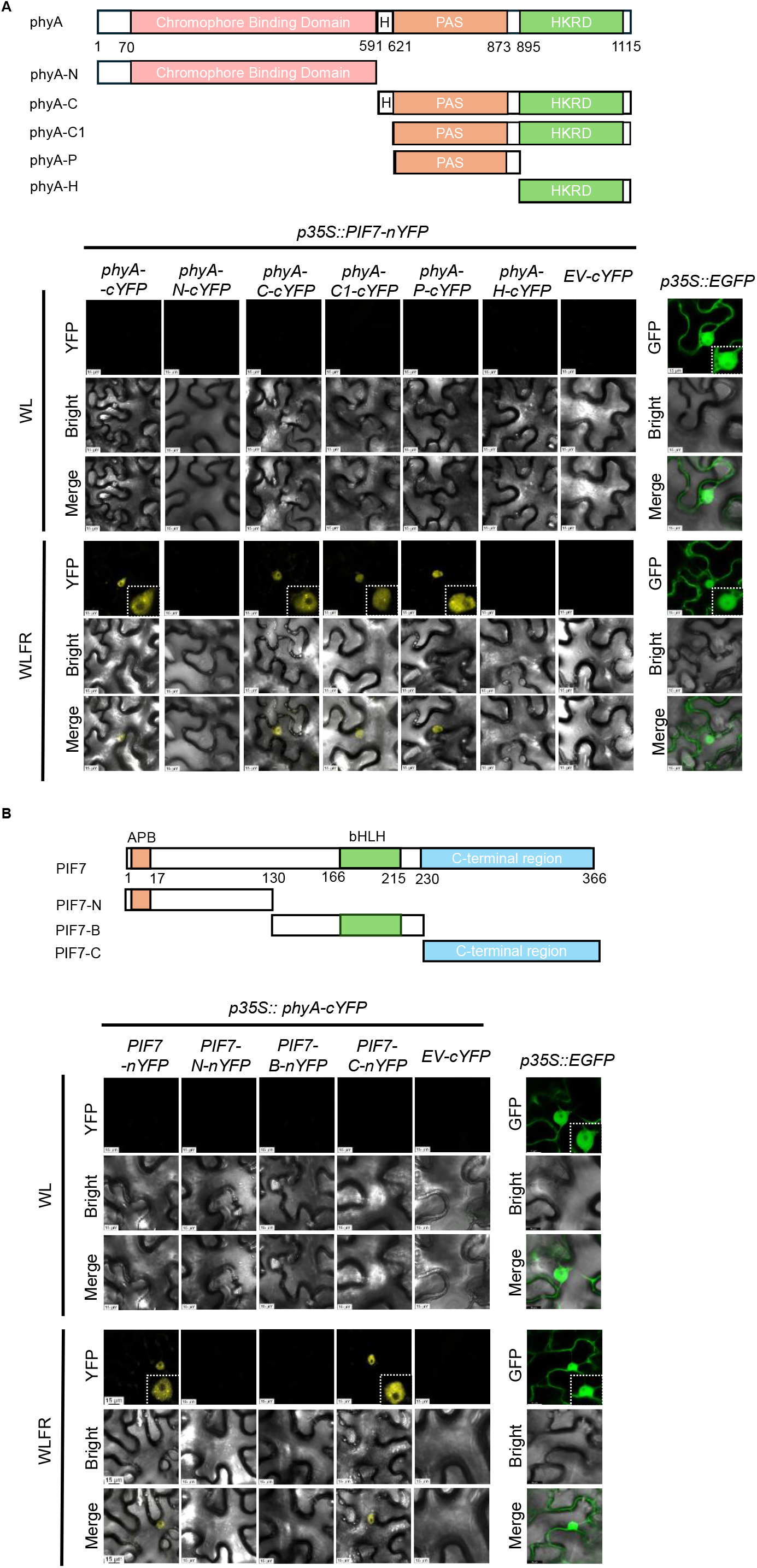
The PIF7-phyA Interaction Depends on the PIF7 C-terminal Region. **(A** and **B)** BIFC assays using truncated constructs identify the PHYA PAS domain and the PIF7 C-terminal Region, as important for the phyA-PIF7 interaction. The construct diagrams depict the complete and truncated PHYA and PIF7 protein structures. Full-length and truncated variants were fused to C-terminal nYFP (PIF7) or cYFP (PHYA) to generate BiFC constructs. The *p35S::EGFP* construct served as a transformation control. PIF7-nYFP + phyA-cYFP was used as a positive control, whereas PIF7-nYFP + *p35S::EV-cYFP* and phyA-cYFP + *p35S::EV-nYFP* served as negative controls. All constructs were transiently expressed in *Nicotiana benthamiana* leaf epidermal cells. 488nm lasers were used to detect the YFP/GFP signals. Representative images from three independent replicates are shown. Scale Bar=15μm.

To identify the region of PIF7 that mediated the interaction with phyA, we generated PIF7-N which contains the ACTIVE PHYTOCHROME BINDING domain [*PIF7-N-nYFP*], PIF7-B, containing the BASIC HELIX-LOOP-HELIX motif [*PIF7-B-nYFP*], and PIF7-C [*PIF7-C-nYFP*] which contains a polyQ-rich, prion-like domain (Fig. 3B and fig. S7). Interestingly we found that only the PIF7 full length control and PIF7-C gave nuclear fluorescence in WLFR. Thus, our data point to the C-terminal region of PIF7 as being important for its interaction with phyA.

### PIF7ox blocks phyA regulation of target genes

To further interrogate the mechanism through which PIF7 inhibits phyA action we initially tested whether PIF7 alters the expression of genes (*CHS, BIN2, FHY3, FHY1, FHL* and *IAA29)* known to be directly regulated by phyA (Fig. 4A) (*32,33*). As determined by qPCR all the genes, except *IAA29*, were upregulated in WT in WLFR conditions, otherwise, the expression patterns fell into two categories. In WLFR conditions, *CHS, BIN2*, and *FHY3* exhibited decreased expression in both *phyA-211* and PIF7ox, while their expression was increased in *pif7-1*. This gene set is upregulated by phyA in WLFR, and their expression is repressed by PIF7. On the other hand, *FHY1, FHYL* and *IAA29* form a group that are upregulated in *phyA-211* and PIF7ox, and for *FHY1* and *FHYL*, expression is lowered in *pif7-1*. These genes are downregulated by phyA in WLFR, and their expression is promoted by PIF7. The common factor for all these genes is that phyA and PIF7 exert opposing control of their regulation, particularly in WLFR. We then sought to establish if the antagonistic action of PIF7 was mediated through direct repression of phyA at target gene promoters. Here we performed chromatin immunoprecipitation (ChIP) using a native phyA antibody, and qPCR using to detect phyA enrichment at *CHS, FHY1*, and *FHL* promoters. For all three genes we found WLFR stimulated phyA enrichment at promoter regions sequences containing G-box motifs (two in *CHS*), but not in G-box-free regions (control, coding region) (Fig. 4, B and C). This enrichment was eliminated in the presence of PIF7ox, suggesting the interaction between PIF7 and phyA is able to block phyA regulation of target genes.

**Fig. 4.**
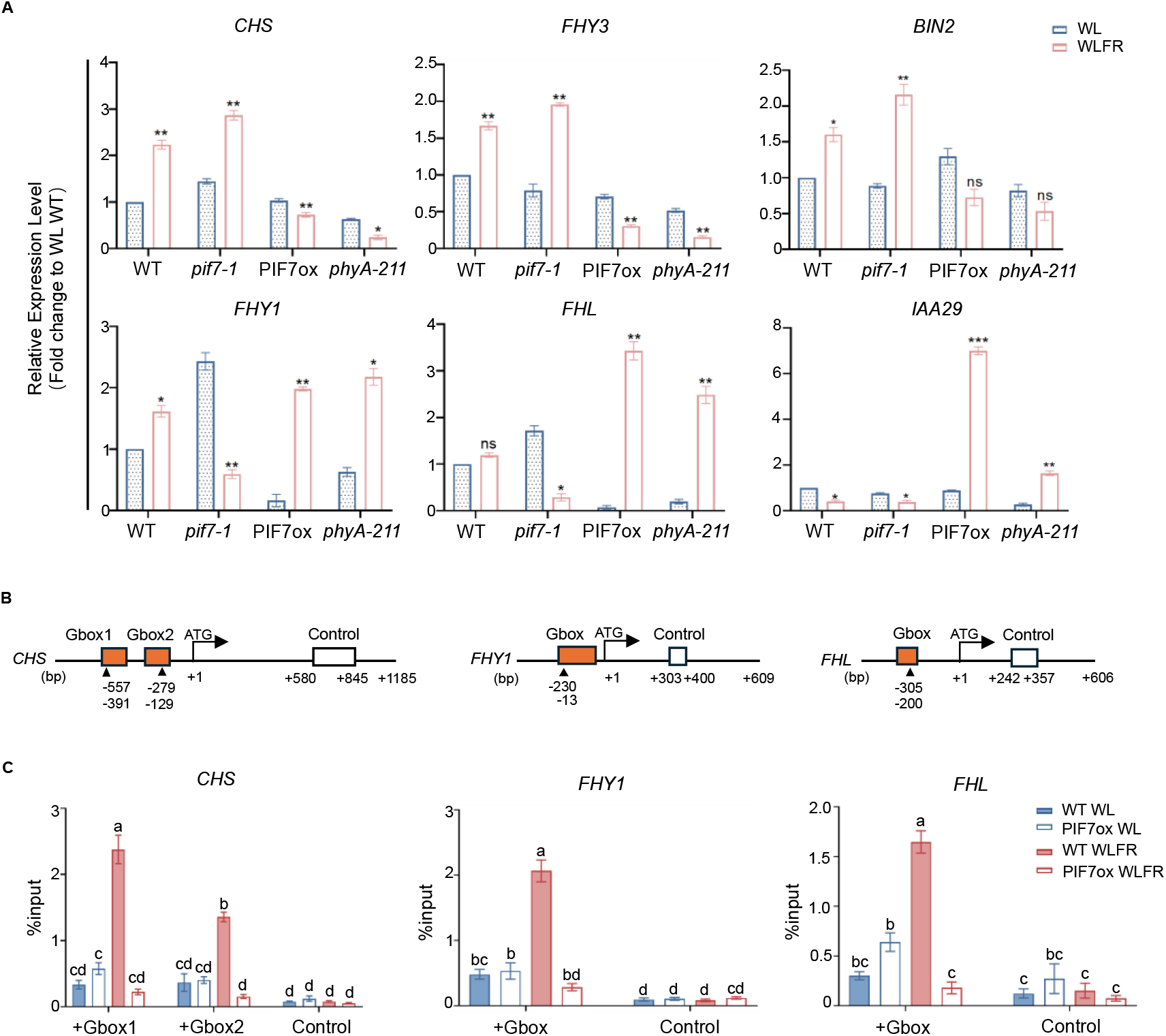
PIF7 inhibits phyA-mediated regulation of target genes. **(A)**Real-time qPCR analysis of *CHS, FHY3, BIN2, FHY1, FHL* and *IAA29* expression in WT (*Col-0*), *pif7-1*, PIF7ox and *phyA-211*. Transcript levels were normalised to the *PP2A* internal control. (**B**) Schematics representations of the *CHS, FHY1* and *FHL* gene and promoter regions The orange boxes indicate primer target regions encompassing G-box elements, while the white boxes denote exon regions used as assay controls. Arrows indicate base pair (bp) position. (**C**) Chromatin immunoprecipitation (ChIP)-qPCR illustrates PIF7ox reduces phyA enrichment at *CHS, FHY1*, and *FHL* promoters under WLFR. Assays were conducted WT (*Col-0*) or PIF7ox seedlings sampled at ZT7 on day 6. Native phyA was immobilised on protein A/G beads using an anti-phyA antibody, and promoter enrichment quantified by Real-time qPCR at G-box promoter regions and coding regions, which served as negative controls. Data shown is normalised to input samples. For (**A** and **C**) data are presented as mean values ± s.e.m., *n*=3 (biological repeats). For (**A**) a Student’s t-test was used to assess statistical significance between the WL/WLFR for each genotype, asterisks indicate significant differences from the WL values (*, P < 0.05; **, P < 0.01; and ***, P < 0.001). For **(B)**a Two-way ANOVA was conducted to assess statistical significance across light conditions for each genotype (α=0.05), followed by Tukey’s HSD post hoc test for pairwise multiple comparisons. Groups that do not differ significantly are indicated by the same letter.

### PIF7 interacts with FHY1 and FHL

In FR-rich conditions FHY1 and FHL play a pivotal role in facilitating the transport of phyA into the nucleus, and they have been linked to the regulation of phyA target genes (*9,10,23*). Notably, FHY1 has been implicated in the recruitment of phyA to the *CHS* promoter through its direct binding to HY5 or PIF3 (*34*). The importance of FHY1 and FHL in our WLFR conditions is evident in the *fhy1-3;fhl-1* double mutant hypocotyl response which is phenotypically similar to *phyA-211* (fig. S8). Consequently, we aimed to determine whether PIF7 interacts with FHY1 and FHL in addition to phyA. Using the *N. benthamiana* transient expression system, we recorded BiFC nuclear interactions between *p35S::PIF7-nYFP (PIF7-nYFP)* and the *p35S::FHY1-cYFP (FHY1-cYFP), p35S::FHL-cYFP (FHL-cYFP)* constructs under WLFR conditions, but not with the *35S::EV-cYFP* control (Fig. 5A. In our split nLUC assay we also observed PIF7-lgbit interactions with FHY1-smbit and FHL-smbit, but not the negative control, under WLFR (Fig. 5B). It is worth noting that in this assay, the interaction signal strength between the shuttle proteins and PIF7 was weaker than that observed between PIF7 and phyA (Fg. 5B). However, we were able to verify these interactions through co-immunoprecipitation using *p35S::FHY1/FHL-nLUC* (FHY1/FHL-nLUC) and *p35S::PIF7-sfGFP* (PIF7-GFP) in *N. benthamiana* (Fig. 5C and fig. S9). As our data shows that PIF7 can interact with phyA, FHY1 and FHL, we postulated that these proteins may form a complex under inductive WLFR conditions. Lending some support for this hypothesis, we found that when FHY1-YFP or FHL-YFP are co-expressed with PIF7-GFP and phyA-RFP in *N. benthamiana* leaves, these constructs localize to the same nuclear regions under WLFR (Fig. 5, D and E).

**Fig. 5.**
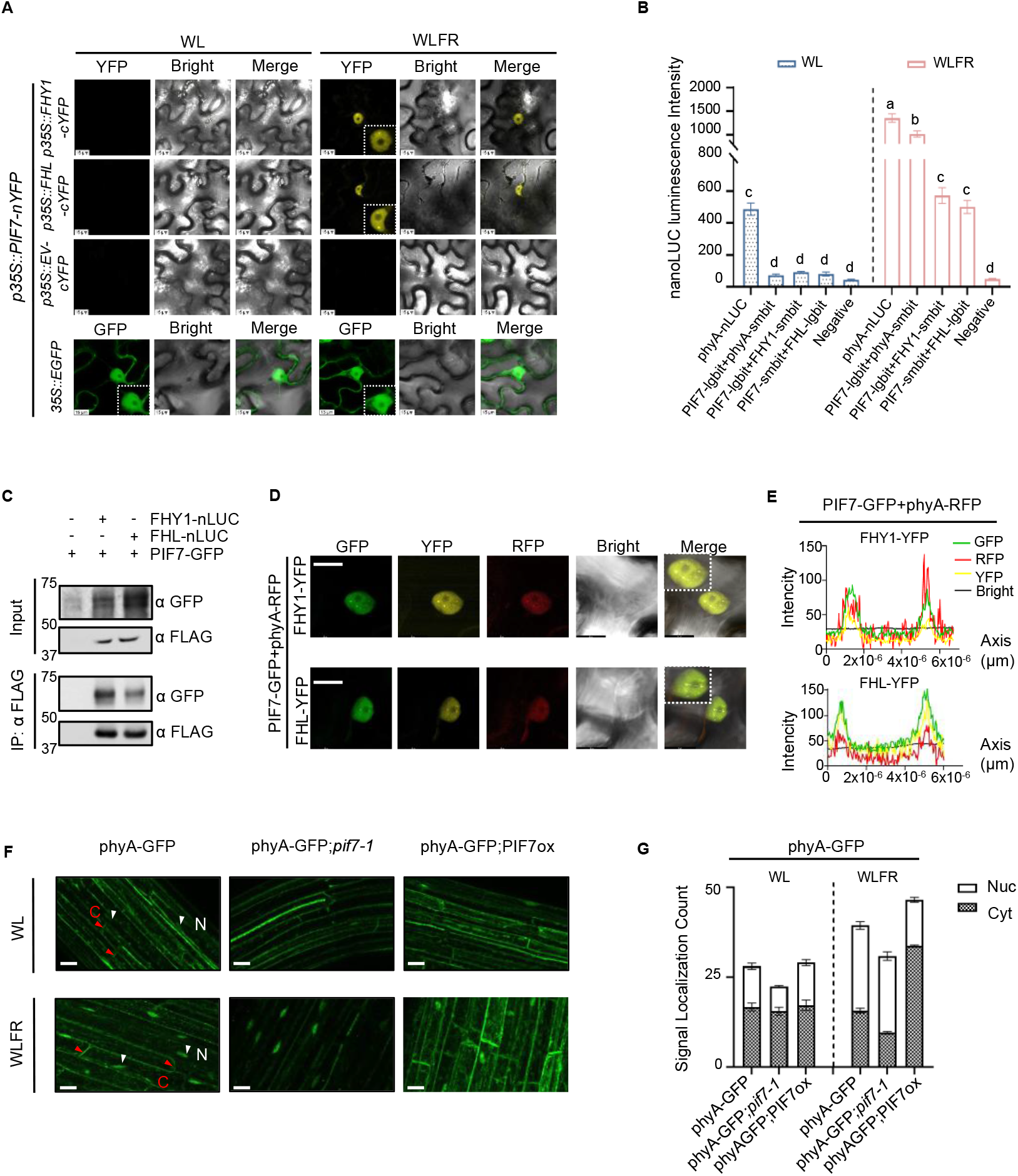
PIF7 interacts with FHY1/FHL in WLFR. (**A**) The BiFC assay illustrates that PIF7 interacts with FHY1 and FHL in WLFR. Interaction fluorescence was detected when *p35S::FHY1-cYFP* (FHY1-cYFP) or *p35S::FHL-cYFP* (FHL-cYFP) were co-expressed with *p35S::PIF7-nYFP* (PIF7-nYFP) in *N. benthamiana* under WLFR. The *p35S::EGFP* used as the transformation control. (**B**) Interactions between PIF7 and FHY1/FHL using a split NanoLUC complementation assay under WL and WLFR. Bioluminescence signals were detected upon co-expression of *p35S::PIF7-lgbitnLUC* (PIF7-lgbit) and *p35S::FHY1-smbitnLUC* (FHY1-smbit) or *p35S::PIF7-smbitnLUC* (PIF7smbit) and *p35S::FHL-lgbitnLUC* (FHL-lgbit) in the transient *N. benthamiana* expression system. The phyA-nLUC and PIF7-lgbit + phyA-smbit served as positive controls in WLFR, whereas PIF7-smbit + phyA-smbit was used as a negative control in WLFR. (**C**) Immunoprecipitation assay demonstrating PIF7 interacts with FHY1/FHL *in vivo* under WLFR. *p35S::FHY1/FHL-nanoLUC × 3FLAG × 10His* (FHY1-nLUC) and *p35S::PIF7-sfGFP* (PIF7-GFP) constructs were transiently expressed in *N. benthamiana* leaves. Anti-FLAG beads were used to pull down FHY1/FHL and its target proteins by Co-IP, while anti GFP antibodies were used to detect PIF7-GFP. (**D** and **E**) Fluorescence assays showing PIF7, phyA and FHY1/FHL co-localise in nuclei and photobodies. *p35S::PIF7-sfGFP* (PIF7-GFP), *p35S::FHY1-YFP* (FHY1-YFP) and *p35S::FHL-YFP* (FHL-YFP), and *p35S::PHYA-RFP* (phyA-RFP) constructs were transiently expressed in *N. benthamiana* leaves. The merged image (D) signifies co-localisation in epidermal cell nuclei, and (E) displays GFP, YFP and RFP channel signals along the white line transect shown in the merged nuclear image. (**F**) Fluorescence imaging to assess the effect of PIF7 on phyA-GFP cellular distribution under WL and WLFR. Lines of phyA-GFP;*pif7-1* and phyA-GFP;PIF7ox were generated by introgressing *pPHYA::PHYA-sGFP* into *pif7-1* and PIF7ox. Images were captured at ZT7 after 6 days of WL or WLFR. The images were collected from mid-hypocotyl region using a 20 × 9.91 objective lens. The white arrows and N label indicate nuclear, while red arrows and C label indicate cytosolic signals. Scale bar = 25 μm. (**G**) Quantification of nuclear vs cytosolic signals from hypocotyl epidermal cells of phyA-GFP, phyA-GFP;*pif7-1* and phyA-GFP;PIF7ox. A total 50 cells were counted in each of 6 biological repeats. For (**G, D** and **F**) 488nm and 552nm lasers were used for detecting GFP/YFP and RFP signals. Scale bar=15μm. For (**B** and **G**) data are presented as mean values ± s.e.m., *n*=12 (leaf numbers) in (**B**) and *n*=6 (biological repeats) in (**G**). In (**B**) a Two-way ANOVA was conducted to assess statistical significance across light conditions for each genotype (α=0.05), followed by Tukey’s HSD post hoc test for pairwise multiple comparisons. Groups that do not differ significantly are indicated by the same letter.

Interestingly, we also found the expression of *ARABIDOPSIS SUMO PROTEASE 1 (ASP1)*, a crucial factor in SUMOylation and the maintenance of FHY1 levels under FR light conditions to be PIF7-dependent (*35*).

In WT, *ASP1* levels are increased by WLFR, a response that is further amplified by *pif7-1* and suppressed by PIF7ox (fig. S10). Thus, *ASP1* may provide a route for PIF7 to indirectly regulate FHY1.

Since PIF7 is able to interact with FHY1 and FHL we were interested to establish if PIF7 affected phyA nuclear shuttling. To address this question, we measured the proportion of cells with nuclear vs cytosolic phyA-GFP signals in the *pif7-1* and PIF7ox backgrounds. In WL, we found this ratio to be similar in all genotypes (Fig. 5, F and G). Whereas in WLFR, we recorded a higher proportion of nuclear phyA-GFP in WT, that was even greater in the *pif7-1* mutant, and diminished in PIF7ox (Fig. 5, F and G). Given that PIF7ox has elevated phyA levels (Fig. 1, E to G), we reasoned that this could potentially skew the proportion of nuclear to cytosolic phyA. However, the observation that *pif7-1* has a slightly higher nuclear fraction of phyA compared to WT, despite having lower phyA levels, suggests that PIF7 may indeed influence the cellular distribution of phyA (Fig. 5, F and G). To verify our results, we performed immunoblot assays that differentiated between nuclear and cytosolic fractions. This method yielded qualitatively similar results showing increased nuclear phyA in *pif7-1* and predominately cytosolic PHYA in PIF7ox under WLFR (fig. S11). Thus, it is possible that PIF7 binding to FHY1 and FHL, may partially disrupt the nuclear transport of phyA.

### phyA-NLS inhibits PIF7ox action

To determine if PIF7 primarily acts by sequestering phyA or whether the PIF7 interactions with FHY1/FHL are also important, we employed *p35S::PHYA-tagRFP:terRbcS* (phyAox) and *pPHYA::PHYA-sGFP-NLS* (phyA-NLS), where phyA is constitutively nuclear (fig. S12) (*30*). In line with earlier findings, under our experimental conditions, phyAox and phyA-NLS restore the elongated *phyA* mutant hypocotyl phenotype to WT length (Fig. 6A). We hypothesized that if PIF7 regulation of phyA was solely due to its interaction with phyA, then PIF7ox should diminish the function of phyAox and phyA-NLS through this mechanism (*30*). However, while this is the case for phyAox, we found phyA-NLS exhibits strong epistasis over PIF7ox for control of hypocotyl length. Similarly, for *CHS* and *IAA29* gene expression under WLFR conditions, PIF7ox suppressed the effects of phyAox, while conversely, phyA-NLS diminished the impact of PIF7ox on *CHS* and *IAA29* (Fig. 6, B and C). These findings are not attributable to variations in phyA levels, as immunoblot analyses show that under WLFR conditions, phyA levels are slightly reduced in phyA-NLS compared to phyAox (fig. S13). Consequently, the potent effects of PIF7ox on phyA action, are unlikely to arise solely from the PIF7-phyA interaction.

**Fig. 6.**
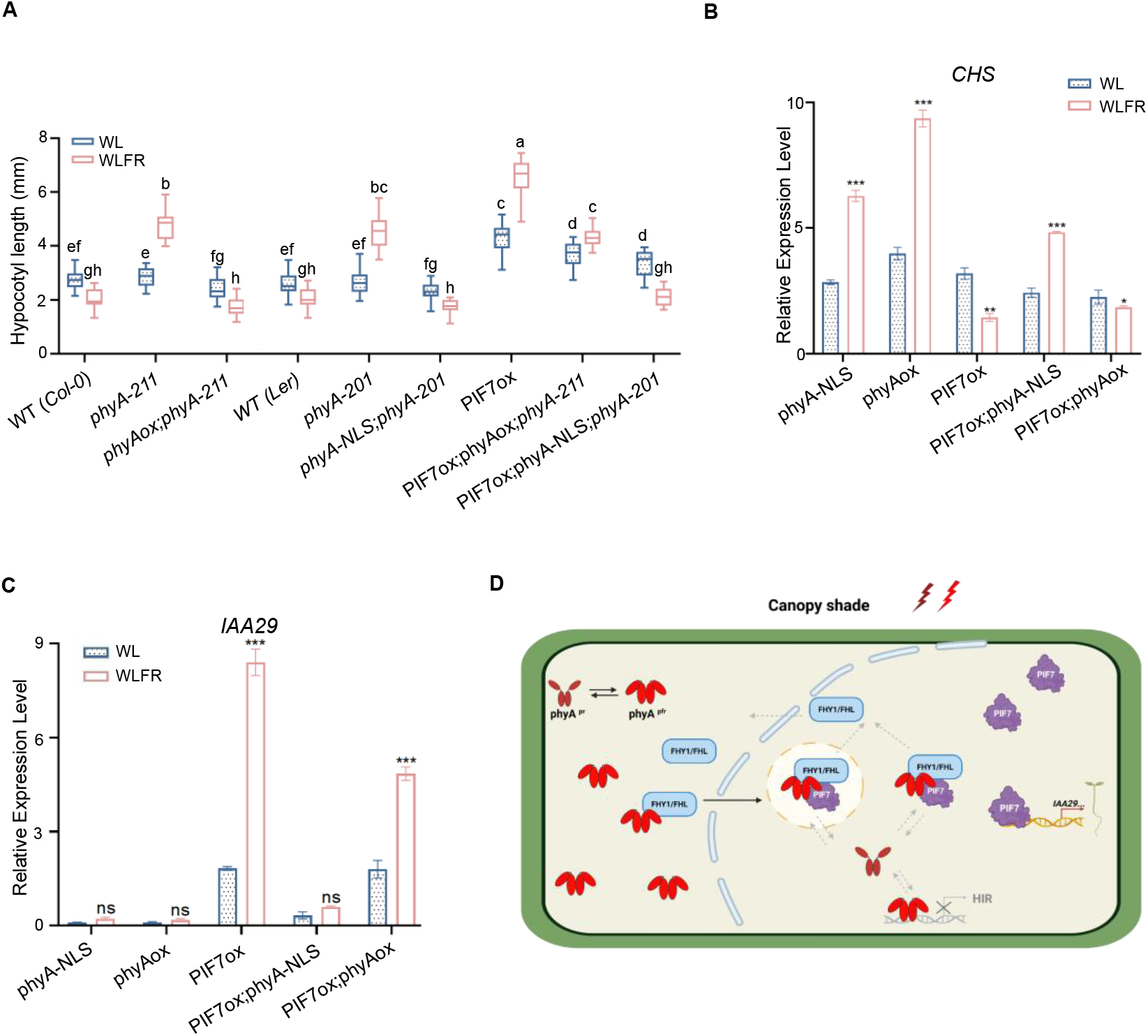
PIF7 cannot suppress phyA-NLS action. (**A**) Hypocotyl length (mm) of 6d-old WT (Col-0), *phyA-211, phyAox;phyA-211*, WT (Ler), *phyA-201, phyA-NLS;phyA-201*, PIF7ox, PIF7ox; *phyAox;phyA-211*, and PIF7ox*;phyA-NLS;phyA-201* seedlings grown in WL or WLFR. Double and triple mutant lines were generated through genetic crossing. (**B** and **C**) phyA-NLS restores phyA control of *CHS* and *IAA29* in PIF7ox seedlings not phyAox. Relative expression of *CHS* and *IAA29* compared to *PP2A* internal control in phyA-NLS*;phyA-201*, (phyA-NLS), phyAox, PIF7ox and PIF7ox;phyA*-*NLS*;phyA-201* (PIF7ox;phyA*-*NLS). Samples were collected at ZT7 on day 6 from seedlings grown in WL or WLFR.(**D**) Schematic depicting the cellular model for PIF7 control of phyA action under canopy shade. In persistent shade conditions phyA is transported to the nucleus by the FHL/FHY1 transporters. PIF7 also accumulates in the nucleus, where it interacts with phyA and with FHL/FHY1 to form photobodies. Here PIF7 exerts a dual effect on phyA by limiting its nuclear translocation and repressing phyA-mediated regulation of target gene expression. Consequently, PIF7 is able to amplify SAS-mediated hypocotyl elongation by blocking the antagonistic action of phyA. The image created in BioRender (https://BioRender.com). For (**A**-**C**), data are presented as mean values ± s.e.m., *n*=24 (seedling numbers) in **a**. *n*=3 (biological repeats) in (**B**) and (**C**). For (**A**) a Two-way ANOVA was conducted to assess statistical significance across light conditions for each genotype (α=0.05), followed by Tukey’s HSD post hoc test for pairwise multiple comparisons. Groups that do not differ significantly are indicated by the same letter. For (**B** and **C**) **a** Student’s t test was used to assess statistical significance between the WL/WLFR for each genotype, asterisks indicate significant differences from the WL values (*, P < 0.05; **, P < 0.01; ***, P < 0.001).

Previous research demonstrated that phyA-NLS completely restores phyA control of hypocotyl growth in *fhy1-3;fhl-1 and phyA-211;fhy1-3;fhl-1*, indicating that phyA-NLS can bypass the requirement for nuclear shuttling (*36*). The effectiveness of PIF7ox on phyAox, which utilizes FHY1/FHL, and not phyA-NLS, suggests that the interaction between PIF7 and FHY1/FHL may be required to suppress phyA action. One possibility is that PIF7 interacts with phyA-FHY1/FHL complexes soon after their nuclear entry and before the complexes disassemble. Interestingly, our fluorescence data illustrate that PIF7-GFP, phyA-RFP and FHY1-YFP or FHY1-YFP co-localize within both the nucleus and photobodies under WLFR conditions (Fig. 5, D and E) (*9*). Collectively, our data indicates that PIF7 forms inactive complexes with phyA and FHY1/FHL, which appear to dampen shuttling and to impede phyA signaling.

## DISCUSSION

PIF7 is known to be a phyB-regulated transcription factor, with a key role in mediating shade-avoidance responses (*18*). This study further reveals that PIF7 is instrumental in controlling phyA signaling, positioning it at the nexus of phyA and phyB pathways. PIF7 operates in persistent shade, conditions that elevate phyA abundance and promote its nuclear accumulation through the FHY1- and FHL-mediated nuclear shuttling mechanism (*9,10*). Our findings demonstrate that PIF7 effectively modulates phyA signaling through direct interactions with phyA as well as with the transport proteins FHY1 and FHL. This regulatory function of PIF7, which occurs within photobodies, directly interferes with phyA-mediated gene expression control.

PIF7 is particularly active in FR-rich shade, which triggers its dephosphorylation, increases its nuclear abundance (fig. S5) (*20*). Since continuous shade also stimulates phyA activity, it creates favorable conditions for interactions between PIF7 and phyA. Indeed, our results demonstrate that PIF7 and phyA physically interact, as evidenced by BiFC (PIF7-nYFP + phyA-cYFP) and split nLUC (PIF7-lgbit and phyA-smbit) transient expression assays, alongside *in vivo* co-immunoprecipitation of PIF7ox and phyA (Fig. 2, B to D). Previously, PIF1 and PIF3 were shown to interact with phyA through their respective Active Phytochrome A-binding (APA) domains (*15,37*). This binding is known to facilitate the proteolysis of PIF1 and PIF3, and to inhibit the activity of PIF3 by isolating it from target promoters (*15,37,38*). PIF7, however, lacks an APA domain, yet it is still capable of binding to phyA. Our data reveal that this interaction is facilitated through the PAS domain of phyA and the C-terminal region of PIF7. Additionally, our ChIP-qPCR assays demonstrate that PIF7 moderates phyA activity by inhibiting its ability to bind to target promoters (Fig. 4, B and C). For the genes *CHS, FHY3*, and *BIN2*, which are activated by phyA in WLFR, and for *FHY1, FHL, IAA29*, which are repressed by phyA, PIF7 counteracts the effects of phyA. Previously, our research demonstrated that PIF7 interacts with and suppresses the transcriptional activity of ANGUSTIFOLIA 3 (AN3), a promoter of leaf cell proliferation (*40*). In this context, PIF7 operates by sequestering and substituting for AN3 at target promoters. These findings suggest that PIF7 can assemble distinct nuclear complexes to modulate the activity of diverse transcriptional regulators.

The proteins FHY1 and FHL are integral to phyA function as they facilitate its nuclear translocation, and FHY1 is proposed to regulate a subset of target genes in concert with phyA (*32,34*). Through the use of BiFc, split nanoLUC and co-immunoprecipitation assays, we have shown that in addition to phyA, PIF7 also interacts with FHY1 and FHL under WLFR (Fig.5, A to C). Under shade conditions, phyA nuclear import is a highly dynamic process that depends on the efficient recycling of FHY1/FHL after dissociation of the nuclear phyA–shuttle protein complex (*8,10, 41,42*). Sequestration of FHY1 and/or FHL by PIF7 could therefore restrict their availability and mobility, thereby reducing their capacity to transport phyA into the nucleus. Consistent with this model, our data reveals an altered cytosolic to nuclear distribution of phyA in hypocotyl epidermal cells of *pif7-1* and PIF7ox (Fig. 5, F and G). Thus, PIF7 appears to affect more than one aspect of phyA function.

In white or red light PIF7 is known to colocalize with phyB in nuclear photobodies (*18,19*). Formation of phyB-PIF7 nuclear condensates is thought to limit the availability of PIF7 for transcriptional regulation (*18,19,43,44*). Conversely, exposure to low R:FR light deactivates phyB, triggering photobody dissolution and releasing PIF7 to bind DNA and mediate chromatin modifications. This study reveals that under persistent shade conditions, PIF7 relocates to distinct photobodies where it co-localizes with phyA and the shuttle proteins FHY1 and FHL (Fig. 5, D and E).

PIF7 is unique amongst PIF proteins as it contains a prion-like domain with a central polyQ stretch of 17 consecutive glutamine residues (fig S7). Prion-like domains typically comprise low-complexity or intrinsically disordered regions that facilitate protein condensation and liquid-liquid phase separation (LLPS). Indeed, PIF7 has been shown to form condensates with LLPS characteristics *in vitro*, which is proposed to facilitate the formation of PIF7-phyB photobodies (*19*). Substituting polyQ with polyA diminishes the capacity of PIF7 to form photobodies, highlighting the importance of polyQ in this process. It is therefore possible that PIF7 also fulfils a similar function in facilitating the formation of PIF7-phyA photobodies. Across organisms, the spatial organization of biomolecules into distinct cellular locations is essential for enabling specialized functions (*45*). Our data suggests that in persistent shade PIF7 may function to sequester phyA along with FHY1 and FHL, thus limiting their operational effectiveness.

We also sought to determine whether the interaction between PIF7 and phyA represents the primary mechanism by which PIF7 regulates phyA signaling. If this were the case, we reasoned that PIF7ox should suppress the effects of both phyAox and phyA-NLS. However, while PIF7ox effectively counteracts the impact of phyAox, it fails to suppress the effects of phyA-NLS on hypocotyl elongation and *CHS* and *IAA29* expression (Fig. 6, A to C). Because phyAox depends on nuclear shuttling for its activity, whereas phyA-NLS is constitutively nuclear and bypasses the FHY1/FHL-mediated transport pathway, these findings suggest that PIF7 interaction with FHY1/FHL, in addition to phyA itself, may be required to attenuate phyA signaling. Therefore, it appears that simultaneously targeting both the receptor and its transport machinery enables PIF7 to effectively suppress this dynamic signaling pathway.

In summary, our study outlines the central role PIF7 plays in regulating phyA activity (Fig. 6 D). Its direct binding to phyA, and the FHY1/FHL transporters, likely represents a rapid, energy-efficient, and highly effective way to stem phyA action. The compartmentalization of phyA and FHY1/FHL into PIF7 photobodies constitutes a dynamic regulatory mechanism that responds to changes in environmental light quality. This subcellular partitioning enables PIF7 to drive hypocotyl elongation while simultaneously reducing the suppressive influence of phyA. Since the modulation of PIF7 by phyB is well-established, our findings position PIF7 at the molecular intersection of phyB and phyA, underscoring its pivotal role in adjusting phyA activity to promote SAS elongation under vegetative shade (*19*).

## METHODS

### Plant material and growth conditions

The wild-type of *Arabidopsis thaliana* seedlings are Columbia-0 (Col-0). The *pif7-1, pif7-2* (*46*), *pif8-1, pif8-2* mutants, and the construction of *pif8-1;phyA-211*, PIF8ox*;phyA-211* and PIF8ox (*15*), PIF7ox-FLASH(*47*), *p35S::GFP-PIF7* (*20*), *p35S::PIF3-LUC* (*48*), *p35S::PIF5-HA, p35S::PIF4-LUC* (*16*), *pPHYA::PHYA-sGFP, pPHYA::PHYA-sGFP-NLS* (*30*), and *pPHYA::PHYA-nanoLUC-3×FLAG-10×His* (phyA-nLUC) (*3*) have been described previously. The *phyA-211* mutant was obtained from NASC collection. *pPIF7::PIF7-nanoLUC-3×FLAG-10×His* (PIF7-nLUC) (Basta) were transformed into the *pif7-1* background. The phyA-GFP-NLS*;pif7-1* and PIF7ox;phyA-GFP-NLS, *pif7-1*;*phyA-211* and PIF7ox;*phyA-211* lines were generated by genetic crossing method. Likewise, the *pif7-1;*phyA-nLUC, PIF7ox;phyA-nLUC, PIF7-nLUC;*pif7-1*;*phyA-211* lines were made through genetic crossing. All the cross lines were verified using PCR or sequencing.

For the hypocotyl growth experiments seeds were first sterilized and then re-suspended in 0.1% (w/v) agar and stored in darkness at 4 ºC for 3 d to stratify. Following this, seeds were individually sown onto plates containing half-strength Murashige and Skoog basal salts mixture (Duchefa Biochemie, Haarlem, Netherlands) dissolved in 1% (w/v) agar (1/2 MS media; pH 5.8). The plates were then transferred to Percival I30BLL growth chambers at 22 ºC (Plant Climatics, Wertingen, Germany). To promote germination seeds were exposed to a 4 h period of white light (WL) (85±5 µmol m^-2^ s^-1^), and then darkness for 20 h, before transferring to experimental photoperiods. For all experiments, seedlings were grown under short day (8L:16D) conditions in either WL or WL with supplementary FR (WLFR) to provide a R:FR ratio of 0.20, PAR=85±5 µmol m^-2^ s^-1^(fig. S1 and Table S1). The condition data collected through L1-180 Spectrometer (LI-COR®).

### RNA extraction and qPCR

For RNA extraction seedlings (n = 36 ± 5 seedling numbers), were harvested at Zeitgeber time 7 (ZT), one hour before dusk. Total RNA was extracted using a phenol-based method (*40*), 300 μL RNA extraction buffer (PH=8.0) was added to each sample, followed by 300 μL 1:1 acidic phenol: chloroform. The supernatant was separated from the pellet by centrifuging for 15 minutes at 13,000 rpm under 4 ºC. Up to 300 μL of the supernatant was transferred to collection tubes with 240 μL isopropanol and 30 μL 3 M sodium acetate. Samples were precipitated at -80 ºC for 15 minutes, then centrifuged for 30 minutes at 13,000 rpm. Pellets were washed with 300 μL of 70% ethanol and then suspended in nuclease-free water and stored at - 80 ºC. Complementary DNA (cDNA) synthesis was performed using the qScript cDNA SuperMix (Quanta Biosciences) as described by the manufacturer. A Lightcycler 480 system (Roche) was employed for qPCR, with 10 μL reactions that incorporated the SYBR Green Master Mix (Roche). The qPCR primers are shown in Supplementary (Table S2). Results were analyzed using Light Cycler 480 (Roche) followed by the ΔΔCt method.

### Hypocotyl measurements and statistical analysis

Images of hypocotyls were taken after 6 days of growth. Hypocotyl length was analyzed using ImageJ (https://imagej.nih.gov/ij/). Data was analyzed by Excel and GraphPad Prism 10. Data are presented as mean values ± s.e.m. A Two-way ANOVA test was used to obtain statistical significance (α=0.05); the Tukey’s HSD posthoc test for multiple comparisons. Levels that are not significantly different are marked with the same letter. Representative images from three independent replicates are shown in all figs.

### Construct assembly

The full-length CDS of *PHYA* and nanoLUC tag (*nanoLUC-3×FLAG-10×His*; NL3F10H) have been described previously (*3*). The CDS of *PIF7* were synthesized by the GeneArt Gene Synthesis service (Invitrogen, MA, USA). The split nanoLUC tags were made by splitting nanoLUC to lgbit (158 aa) and smbit (11 aa). Mobius Assembly was used to insert the constructs into a mUAV plasmid (L0, CamR) with *AraI* restriction site and plasmid assembly was performed and subsequently verified by the Edinburgh Genome Foundry (*49*). The full-length CDS of *FHY1* and *FHL* parts were amplified with Q5 High-Fidelity DNA polymerase (New England Biolabs, MA, USA) from the Col-0 cDNA by gene-specific primers. The *PHYA-C, PHYA-C1, PHYA-P, PHYA-H, PIF7-N, PIF7-B* and *PIF7-C* part cloned on the CDS region of *PHYA* and *PIF7* (L0 parts). These parts were assembled into a mUAV plasmid and sequenced by Source Bioscience (Cambridge, UK). All L0 parts will assemble in pMAP B L1 (KamR) vector with *BsaI* restriction site. All the constructs for the tobacco (*Nicotiana benthamiana*) assay used a CaMV35S promoter, which is synthesized by the GeneArt Gene Synthesis service (Invitrogen, MA, USA).

To construct *pEXP601-pPIF7::PIF7-nanoLUC-3×FLAG-10×His*, a 2740 bp genomic fragment was generated, encompassing the PIF7 promoter region (1015 bp), 5’UTR (161 bp), and 1564 bp ORF, excluding the stop codon of exon 6 of the AT5G61270.1 gene model. This fragment was PCR-amplified using specific primers with Phusion High-Fidelity DNA Polymerase (NEB, M0530S). The resulting amplicon was subsequently recombined into pDONR221 using BP II clonase (Invitrogen, 12535029), and its sequence verified by Sanger sequencing (Genepool, Edinburgh Genomics, Edinburgh, UK). The genomic regions were then recombined into *pGWB601:nanoLUC-3×FLAG-10×Hiss* using LR Clonase (Invitrogen) (*50*). The final construct was transformed into Agrobacterium AGL1. Primers used for construct assembly are listed in Supplementary (Table S3).

### Transient Expression Assay

For this assay, constructs were transferred into the Agrobacterium AGL1 strain and cultured at 28 ºC for two days, before infiltration into 4-week-old *Nicotiana benthamiana* leaves with P19. The transient expression solution contained 10 mM MES buffer (pH 5.6), 10 mM MgCl_2_ and 200 mM aceto-syringone. Following infiltration plants were grown for 3 days in short day (8L:16D, at 22 ºC) conditions in either WL or WLFR to provide a R:FR ratio of 0.20. Samples were analyzed at ZT7 on day 3.

### Confocal Imaging, Colocalization, and BiFC

Images of GFP, YFP and RFP fluorescence were obtained with a Leica confocal micro-scope (Leica SP8) using 488 nm and 552 nm lasers. For BiFC and co-localization assay all images were taken at ZT7 on day 3 after WL/WLFR treatments. The co-localization signals were analyzed by LAS X Office 1.4.7.2982 and plotted by GraphPad Prism 10. *pPHYA::PHYA-sGFP(NLS)*, phyA-GFP;*pif7-1*, phyA-GFP;PIF7ox and *p35S::GFP-PIF7* transgenic lines images were taken at ZT7 within 30 minutes after 6 days of WL/WLFR treatments. The Nuclear and cytosolic signals were quantified from 50 cells in the mid-hypocotyl region within a 20 × 1.36 objective lens. Nucleus and cytoplasmic signals were quantified after eliminating background interference by Image J. Then plotted in GraphPad Prism 10 (*32*). Imaging experiments were comprised of three biological replicates, with 6 seedlings per replicate.

### Immunoblot Analysis and Co-immunoprecipitation Assays

Each sample comprised 0.5 g of Arabidopsis seedling tissue, harvested on day 6 at ZT7. The seedlings grounded into fine powder in liquid nitrogen and then homogenized in double the volume of pre-cooled lysis buffer which contained 1× Protease inhibitor (Sigma-Aldrich, P9599) and 1× HALT-Phosphatase inhibitor (Thermo Scientific) (125 mM Tris-HCl (pH 7.5); 0.25mM EDTA (pH 7.5), 150 mM NaCl, 5% glycerol, 0.1% Triton X-100, 0.5% NP-40). Then mixture samples were centrifuged at 12,000 rpm for 10 mins at 4 ºC and then transferred to lysate mix with 1× SDS loading buffer (containing 0.1 M DTT) at a 1:1 volume ratio. Samples were incubated at 80 ºC for 10 mins, followed by a centrifugation step, before western blotting.

For Co-immunoprecipitation, the different light-treated Arabidopsis seedlings(0.5-1g) or *Nicotiana benthamiana* leaves (2-3 g) were ground to a fine powder in liquid nitrogen and then homogenized in double the volume of pre-cooled lysis buffer which contained 1× Protease inhibitor (Sigma-Aldrich, P9599) and 1× HALT-Phosphatase inhibitor (Thermo Scientific) (125 mM Tris-HCl (pH 7.5); 0.25mM EDTA (pH 7.5), 150 mM NaCl, 5% glycerol, 0.1% Triton X-100, 0.5% NP-40) after lysis we kept this mixture on ice for 30 minutes and centrifuge 10 minutes with 12,000 rpm. The supernate was filtered twice through two layers of Mira cloth (Merck Millipore). After 100 ul supernatant, consisting of total protein extracts as input, and the other supernatant were incubated with Protein A/G beads (MCE Magnetic Agarose) for 1 h at 4 °C to remove adventitious binding proteins. After incubating with Protein A/G beads, moved beads from supernatant by Magnetic Separation Rack. Then anti-FLAG/His beads (Anti-FLAG M2 agarose gel Sigma-Aldrich, Cat.No.

A2220; Anti-His Magnetic Beads, Cat. 209, MCE) were added into treated protein extracts and kept overnight at 4 ºC (Before adding the antibody-beads, used wash buffer wash beads 3 times). The antibody/protein complex was washed with wash buffer (125 mM Tris-HCl [pH 7.5]; 0.25mM EDTA [pH 7.5], 150 mM NaCl, 5% glycerol, 0.1 % Triton X-100) 5 times. Then, 50 μL of 1× SDS loading buffer were added to each sample and incubated at 80 ºC for 8 mins.

To detect native phyA protein, an anti-Phytochrome A antibody (PHYTOAB, PHY1907) was used at 1:1,000. To detect the PIF7ox-FLASH and *p35S::FHY1/FHL-nanoLUC × 3FLAG× 10His* an anti-FLAG M2 antibody (Sigma-Aldrich, F1804) was used at 1:2,000. To detect the *p35S::PIF7-sfGFP* an anti-GFP anti-body (Roche, ab290) was used at 1:3,000. Anti-RPN6 antibody (Agrisera, AS121858) was used at 1:3,000 as internal control. Secondary antibodies (1:3,000, anti-mouse IgG (cell signaling, 7076s); 1:4,000 Goat anti-rabbit-IgG (Abcam, AB6013) were used for the ECL Prime Western Blotting System. The signals were detected by Azure 300 Imager with ECL prime western blotting detection reagents (Cytiva, RPN2232).

### NanoLUC and split nanoLUC assays

NanoLUC bioluminescence assay was conducted as previously described (*3*). Single Arabidopsis seedlings were grown in individual wells, within a 96-well plate, each containing 200 µl 1/2 MS media. Plates were sealed with cling film and placed under 8L:16D (WL) for 6 days. Before imaging, 50 µl of 1:50 water dilution of Nano-Glo buffer (Promega, Nano-Glo®Luciferase Assay) was added to the seedlings in each well. Plates were sealed and moved to experimental light conditions. Bioluminescence was recorded every hour (for 1.5 s) for several days using the TriStar2 LB942 multimode microplate reader (Berthold Technologies, Germany). In Fig. 1G, the data were analyzed by averaging individual seedling measurements and then calculating the fold change relative to the ZT0 data for each biological replicate. Subsequently, the results were plotted using GraphPad Prism 10. A total of 9 biological replicates were conducted, with each replicate comprising at least 6 seedlings. In fig. S4, the data was analyzed by BioDare2 software, applying amp&baseline detrending and normalization steps, before plotting in GraphPad Prism 10 (*51*)

The split nanoLUC assay utilized the *N. benthamiana* transient expression system described above. Following WL/WLFR treatments for 3 days, 1cm x 1cm leaf samples (*n* =12) were transferred to 96 wells plates containing 100 µl diluted (1:50) Nano-Glo for imaging with a plate reader. The data analyzed by Excel and GraphPad Prism 10. Experiments repeated at least 3 times.

### Chromatin Immunoprecipitation (ChIP)-qPCR

We employed ChIP-qPCR methods described previously (*52*). For each sample 0.5-1 g of tissue was extracted from 6-day old seedlings at ZT7. Cross-linking was induced with 1% formaldehyde under vacuum infiltration for 15 mins (up to -70 kPa), then 2.5 ml of 2 M glycine was added to quench crosslinking. DNA was fragmented into an average size of ∼150 bp in Diagenode BioRuptor Plus, (30 cycles of high power 30s ON / 30s OFF). Anti-PHYA antibody (PHYTOAB, PHY1907,1:300) incubated with protein A/G beads (MCE Magnetic Agarose) in 4 ºC overnight. And the following day, samples were incubated with protein A/G bead/PHYA antibody complex for 6 hours at 4 ºC. DNA was extracted using phenol/chloroform (1:1, vol/vol). Protein enrichment at DNA regions was quantified using qPCR with the primers detailed in **Supplementary Table 2**. Enrichment was determined by %Input method (%Input=2 ^ [CtInput-CtChIP] × IDF × 100%).

### Nuclear-Cytoplasmic Fractionation

Nuclear fractionation was performed essentially as previously described with the following modifications (*53*). Briefly, 0.5-1 g of Arabidopsis seedlings ground into fine powder in liquid nitrogen and then homogenized in double the volume of pre-cooled (4 °C) lysis buffer (20 mM Tris-HCl, pH 7.4, 25% Glycerol, 20 mM KCl, 2 mM EDTA, 2.5 mM MgCl_2_, 250 mM Sucrose and 5 mM DTT, add 1× protease inhibitor cocktail (Roche) and 1× HALT-Phosphatase inhibitor added in fresh). The sample was filtered twice through two layers of Mira cloth (Merck Millipore). The flow-through was spun 10 min at 2000 g, 4 °C. Supernatant, consisting of the cytoplasmic fraction, was centrifuged 20 min at 13,000 g, 4 °C and collected. The pellet, containing the nuclear fraction, was washed six times with 2.5 mL of nuclear resuspension buffer NRBT1 (20 mM Tris-HCl, pH 7.4, 25% Glycerol, 2.5 mM MgCl_2_, 0.2% Triton X-100) and centrifuged 3 min at 1,500 g, 4 °C. The pellet was resuspended in 300 µL of pre-cooled NRBT2 (250 mM Sucrose, 10 mM Tris-HCl, pH 7.5, 10 mM MgCl_2_, 1% Triton X-100, 0.035% β-mercaptoethanol, and 1× protease inhibitor cocktail added in fresh), and carefully overlaid on top of 300 µL NRBT3(1.7 M Sucrose, 10 mM Tris-HCl, pH 7.5, 2 mM MgCl_2_, 0.15% Triton X-100, 0.035% β-mercaptoethanol, and 1× protease inhibitor cocktail added in fresh). These were centrifuged at 16,000 g for 55 min at 4 °C, and the final nuclear pellet was resuspended in 2× SDS loading buffer. As quality controls for the fractionation, RbCL (Agrisera, AS03037, 1:3,000) protein was used as the cytoplasmic control, and histone H3 (Abcam, ab18521, 1:3,000) was used as the nuclear control in immunoblot assays.

### Statistics and reproducibility

All measurements were performed with three or more independent replicates from separate experiments. The exact sample size and statistical test for each experiment are described in the relevant figure legends. All results are presented as the means ± SEM. Statistical analyses were conducted using GraphPad Prism 10.

The Student’s t test was used to assess statistical significance in Real-time q-PCR data. The other statistical significance used One-way ANOVA test (α=0.05) with the Dunnett test and Two-way ANOVA (α=0.05) with Tukey’s HSD posthoc test. The Levels that are not significantly different are marked with the same letter.

## ACKNOWLEDGEMENTS

This research was supported by University of Edinburgh Darwin Trust Scholarship awarded to MK. Z and the Biotechnology and Biological Sciences Research Council (BB/N005147/1), and Leverhulme Trust (RPG-2024-082) grants awarded to K.J.H. We thank Dr. Rist Van de Weyer who generated PIF7ox;*phyA-211* and first observed the PIF7ox hypocotyl phenotype in WLFR.

We thank Prof. Lin Li, Prof. Dr. Andreas Hiltbrunner, Dr. Gabriela Toledo-Ortiz and Dr. Mathias Zeidler who kindly shared *p35S::GFP-PIF7, p35S::PHYA-RFP, pPHYA::PHYA-sGFP* (NLS) and *fhy1-3;fhl-1* seeds.Thank Prof. Giltsu Choi for sharing *pif8-1;phyA-211, PIF8ox;phyA-211* and PIF8ox lines.

## AUTHOR CONTRIBUTIONS

MK.Z and K.J.H. planned and designed the research. MK.Z conducted the experiments, which were analyzed by K.J.H. and MK.Z. A.R. generated the *pPIF7::PIF7-nanoLUC* line. The manuscript was prepared by MK.Z and K.J.H, subsequent editing performed by K.J.H and A.R.

