## Supplemental files for "PHYTOCHROME INTERACTING FACTOR 7 moderates the activity of phytochrome A in canopy shade conditions"

Science Advances
AAAS

### Supplementary Materials for

#### PHYTOCHROME INTERACTING FACTOR 7 moderates the activity of phytochrome A in canopy shade conditions

Mengke Zhou *et al.*

**This PDF file includes:**

Figs. S1 to S13
Tables S1 to S3

**Fig. S1.**

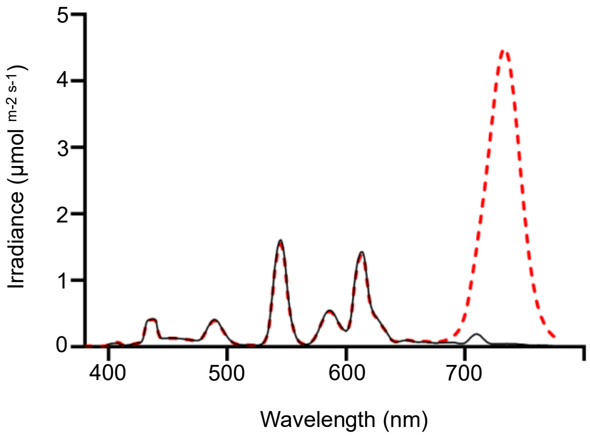


**Fig. S1. Spectral outputs of cabinets used in experimental work.** Light spectra in Percival I30-BL incubators fitted with Luxline Plus F18W/840 fluorescent tubes. The solid black line shows emission of fluorescent tubes (i.e. WL or high R:FR), and the dashed red line shows the spectrum with added FR from OLSON 150 6+ Series FR LED strips, which generate a low R:FR of 0.2 for the WLFR regime.

**Fig. S2.**

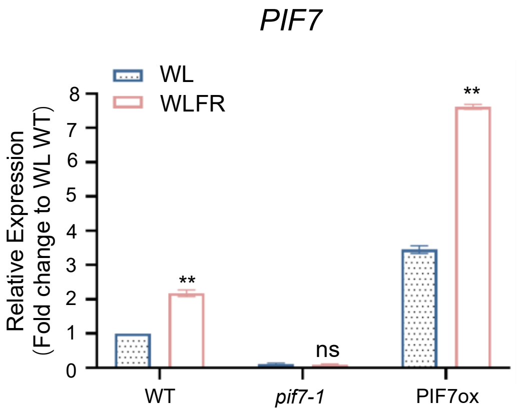


**Fig. S2. *PIF7* mRNA levels in PIF7ox.** Expression of *PIF7* mRNA levels relative to the *PP2A* internal control, determined by real-time qPCR. Samples were collected at ZT7 (1 hour before dusk) on day six, from seedlings grown in WL or WLFR. Data are presented as mean values ± s.e.m., with *n*=3 (biological repeats). Asterisks indicate significant differences from the WL values (*, P < 0.05; **, P < 0.01; and ***, P < 0.001; Student's t test).

**Fig. S3.**

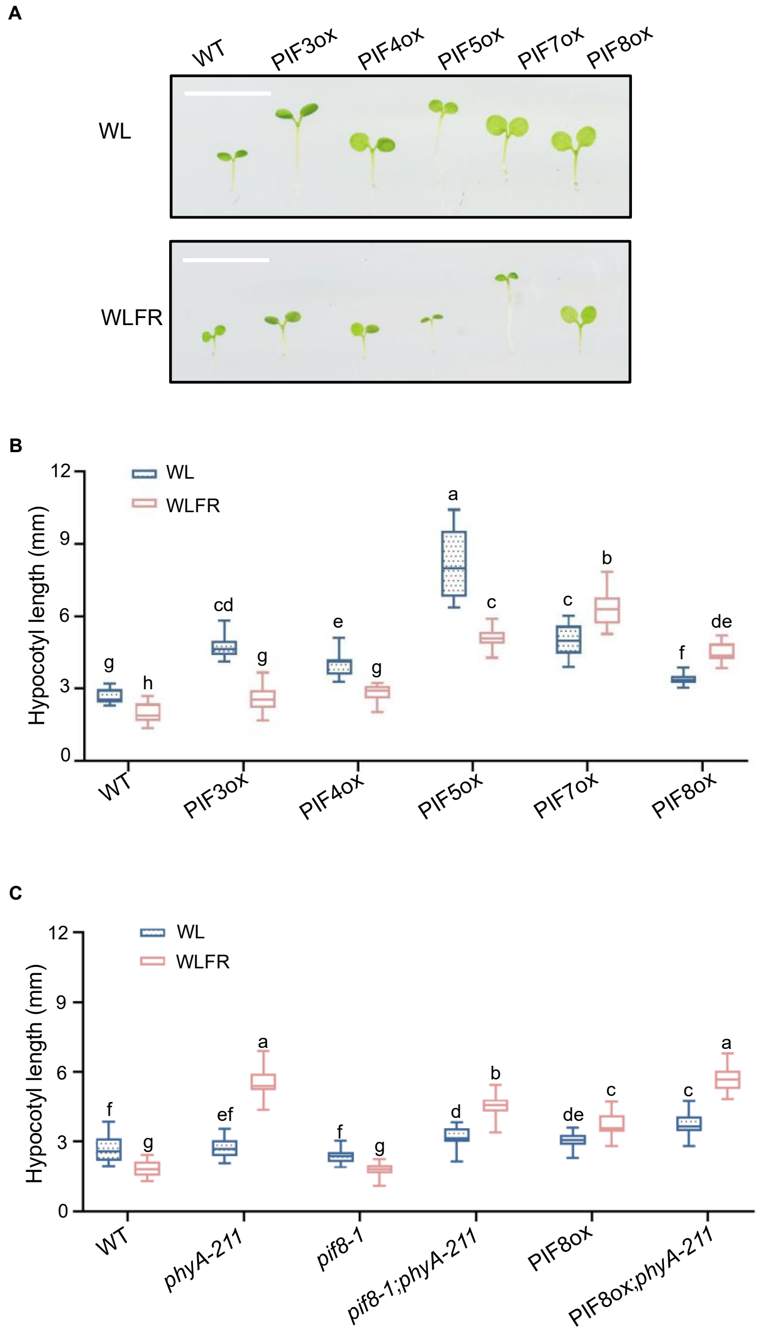


**Fig. S3. Hypocotyl length of PIF lines in WL and WLFR. (A and B)** Images and hypocotyl length (mm) of WT (*Col-0*)**,** *p35S::PIF3-LUC* (PIF3ox), *p35S::PIF4-LUC* (PIF4ox), *p35S::PIF5-HA* (PIF5ox), *p35S::PIF7OX-FLASH* (PIF7ox) and *p35S::MYC-PIF8OX*(PIF8ox) seedlings. In

**(A)** the scale bar =1 cm. **(C)** Hypocotyl length of WT (*Col-0)*, *phyA-211*, *pif8-1*, *pif8;phyA-211*, PIF8ox and PIF8ox;*phyA-211* seedlings. Lines were generated as previously described (*15,16,40,41*) .

Seedlings were grown for 6 days 8 h light (PAR = 85 ± 5 μmol $m^{-2}$ $s^{-1}$), 16 h dark photocycles (WL) or with supplementary FR (R:FR 0.20) (WLFR) at 22°C. Hypocotyl length was measured using ImageJ. Data are presented as mean values ± s.e.m., *n*=27 in **(B)**; *n*=28 in **(C)** (seedling numbers). A Two-way ANOVA was conducted to assess statistical significance across light conditions for each genotype (α=0.05), followed by Tukey's HSD post hoc test for pairwise multiple comparisons. Groups that do not differ significantly are indicated by the same letter.

**Fig. S4.**

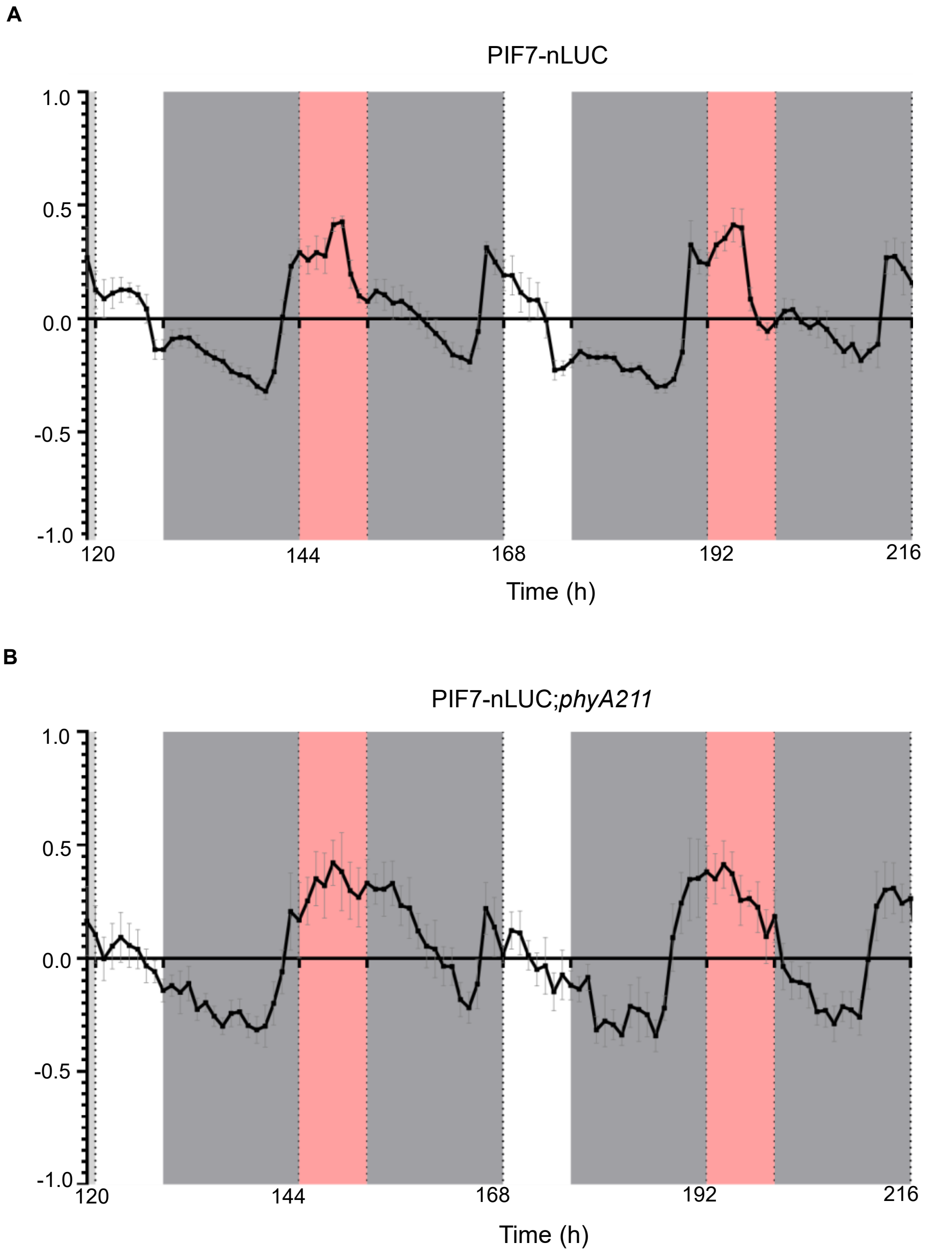


**Fig. S4. Diel rhythms of PIF7-nLUC.** Bioluminescence traces are shown for *pPIF7::PIF7-nanoLUC;pif7-1* (PIF7-nLUC), and *pPIF7::PIF7-nanoLUC;pif7-1;phyA-211* (PIF7-nLUC;*phyA-211*) lines. Seedlings were grown for 6d in 8hL:16hD (PAR = 85 ± 5 μmol $m^{-2}$ $s^{-1}$) before imaging over 3 days. White light (WL) periods are shown in white, while low R:FR (0.20) periods are shown in red, and grey represents darkness. The raw data were analyzed using BioDare2 (*51*), and data are presented as mean values ± s.e.m, n=5 (seedling numbers). Representative data from three independent replicates are shown.

**Fig. S5.**

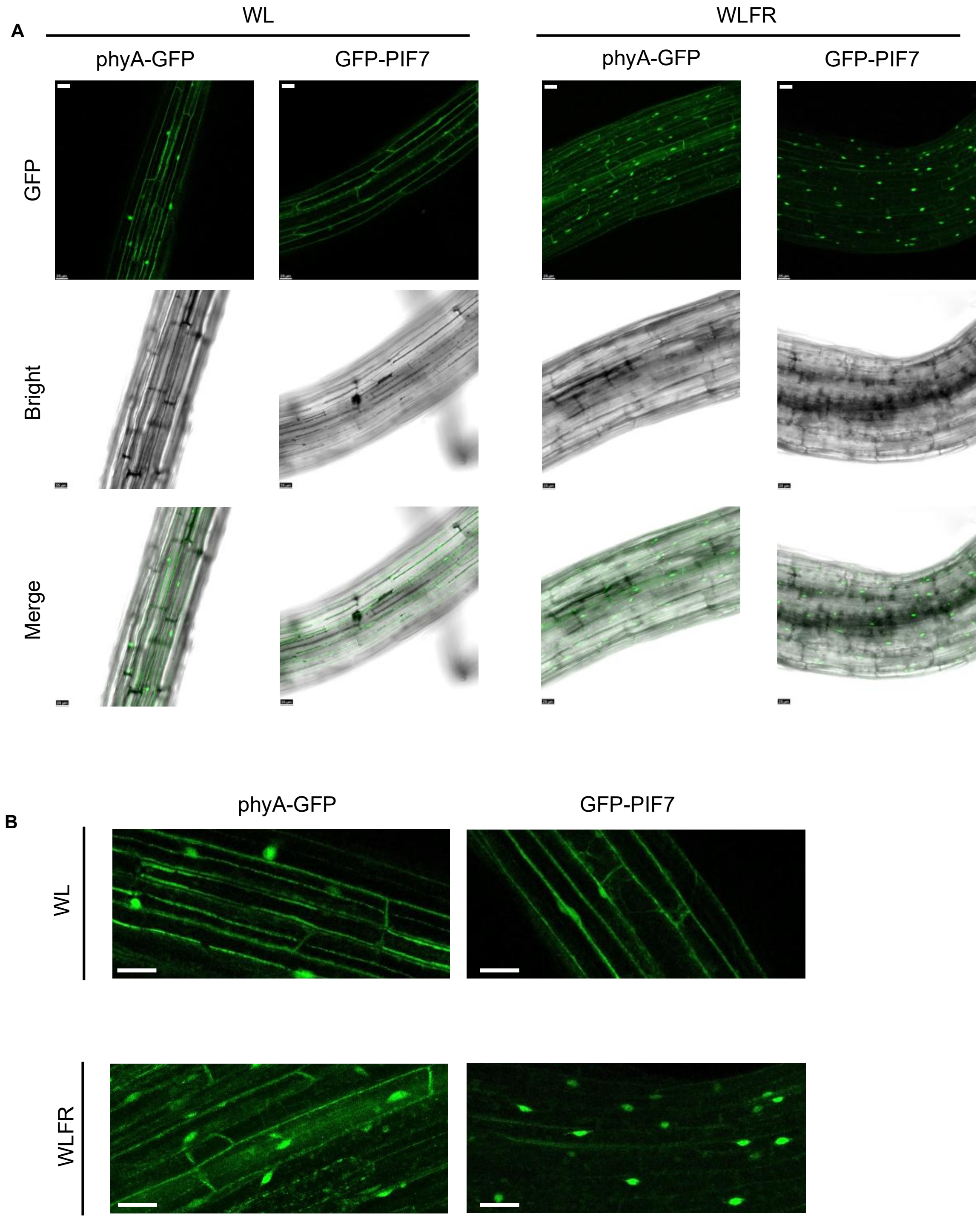


**Fig. S5. PIF7 and phyA both accumulate in nuclei under persistent shade. (A-B)** Cellular distribution of *pPHYA::PHYA-sGFP* (phyA-GFP) and *p35S::GFP-PIF7* (GFP-PIF7) in Arabidopsis hypocotyl epidermal cells. Seedlings expressing phyA-GFP or GFP-PIF7 were grown in WLFR or WL for 6 days. GFP fluorescence signals were detected over 30 minutes using a 488 nm laser. The images were collected from mid-hypocotyl region using a 20 × 1.36 objective lens. Scale bar = 25 μm. Representative images from three independent replicates are shown.

**Fig. S6.**

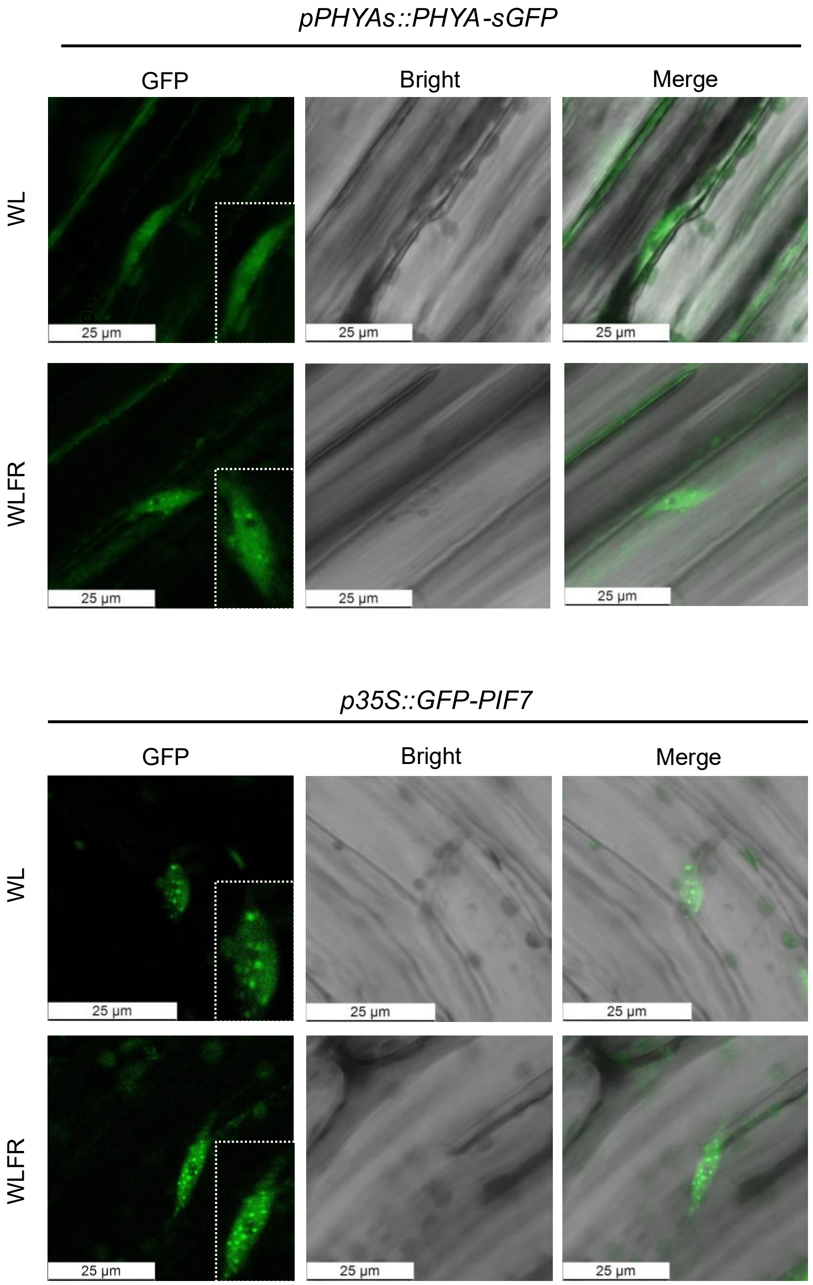


**Fig. S6. PIF7 and phyA both form nuclear photobodies in persistent shade. (A** and **B)** Images were captured at ZT7 on day 6 from transgenic lines expressing *pPHYA::PHYA-sGFP* (phyA-GFP) and *p35S::GFP-PIF7* (GFP-PIF7) were taken at ZT7, grown under WL or WLFR conditions. GFP signals were captured over a 30-minute period, using a 488 nm laser. The images were collected from mid-hypocotyl region using a 20 × 9.91 objective lens. Scale bar = 25 µm. Representative images from three independent replicates are shown.

**Fig. S7.**

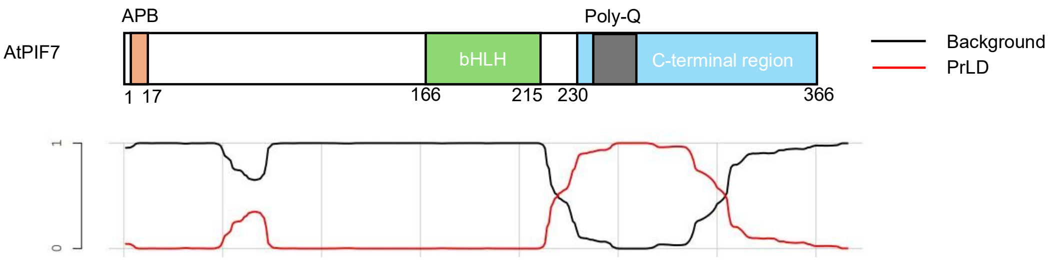


**Fig. S7. Prion-Like Amino Acid Composition Prediction.** Diagram of the PIF7 protein showing motif locations. The PLAAC tool: 'PLAAC: Finding Sequences with Prion-Like Amino Acid Composition' from MIT (using *Arabidopsis thaliana* normalization) predicts Prion-Like Domain (PrLD) in the proximal section of the C-terminus, aligning with the poly-Q region. The red line indicates the PrLD prediction scores (ranging from 0 to 1).

**Fig. S8.**

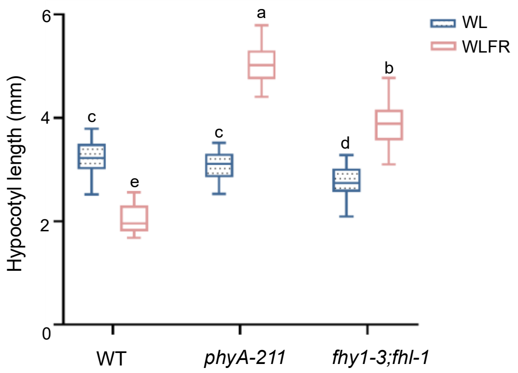


**Fig. S8. The *fhy1-3;fhl-1* is phenotypically similar to *phyA-211* in WLFR.** Hypocotyl length (mm) of WT (*Col-0*), *phyA-211* and the *fhy1-3;fhl-1* double mutant grown for 6d under WL and WLFR (*54*). Data are presented as mean values ± s.e.m., *n*=25 (seedling numbers). a Two-way ANOVA was conducted to assess statistical significance across light conditions for each genotype (α=0.05), followed by Tukey's HSD post hoc test for pairwise multiple comparisons. Groups that do not differ significantly are indicated by the same letter.

**Fig. S9.**

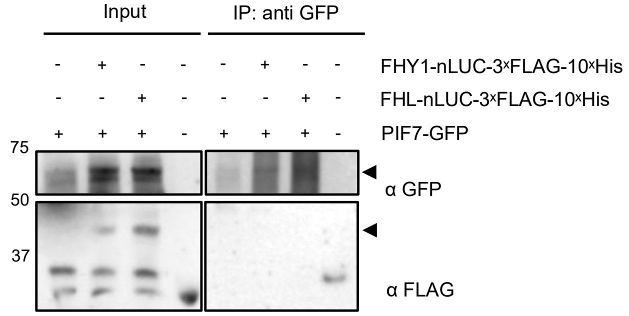


**Fig. S9. PIF7 does not interact with FHY1 or FHL under WL conditions**. Immunoprecipitation assay using *p35S::FHY1/FHL-nanoLUC×3FLAG×10His* and *p35S::PIF7-sfGFP* constructs transiently expressed in *N. benthamiana* leaves under WL. Anti-GFP beads were used to pull down and to detect PIF7-GFP, and an anti-FLAG antibody was used to detect FHY1/FHL-nLUC. We were only able to detect FHY1/FHL-nLUC in Input, suggesting PIF7 and FHY1/FHL do not interact in WL.

**Fig. S10.**

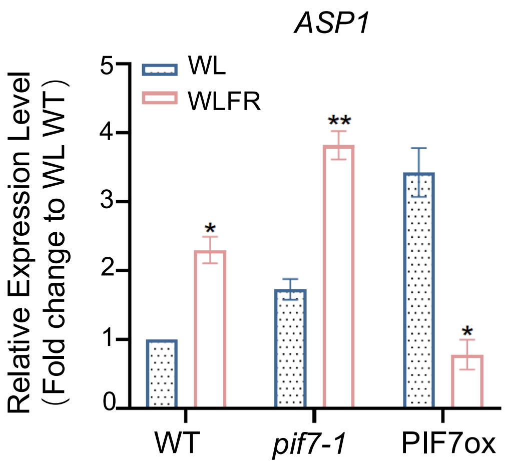


**Fig. S10. PIF7 reduces *ASP1* mRNA levels under WLFR.** Real-Time qPCR analysis of *ASP1* expression in WT, *pif7-1* and PIF7ox, *PP2A* was used as internal control. Data are presented as mean values ± s.e.m., *n*=3 (biological repeats). The Student's t test was used to assess statistical significance between the WL/WLFR for each genotype, asterisks indicate significant differences from the WL values (*, $P < 0.05$; **, $P < 0.01$). Representative data from three independent replicates are shown.

**Fig. S11.**

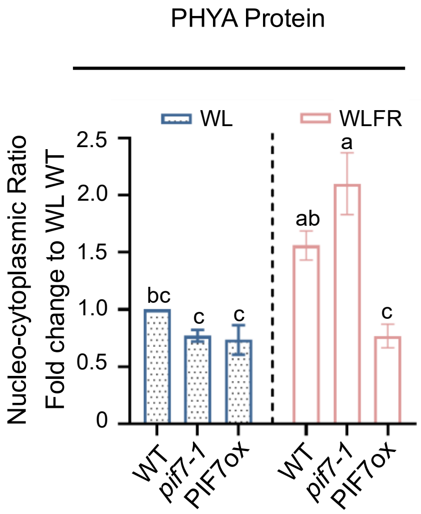


**Fig. S11. PIF7ox enhances PHYA protein accumulation in the cytosol under WLFR conditions.** Six-day-old seedlings were sampled as ZT7. Immunoblot assays were conducted using an anti-PHYA antibody, using anti-RbCL (Rubisco) as the cytosolic control, and anti-Histone 3 (H3) as the nuclear control. The cytoplasmic and nuclear phyA was normalised to RbCL and H3, respectively. The N/C ratio for each genotype is shown relative to WT (*Col-0*), WL control. Data are presented as mean values ± s.e.m., n=3 (biological repeats). A Two-way ANOVA was conducted to assess statistical significance across light conditions for each genotype (α=0.05), followed by Tukey's HSD post hoc test for pairwise multiple comparisons. Groups that do not differ significantly are indicated by the same letter.

**Fig. S12.**

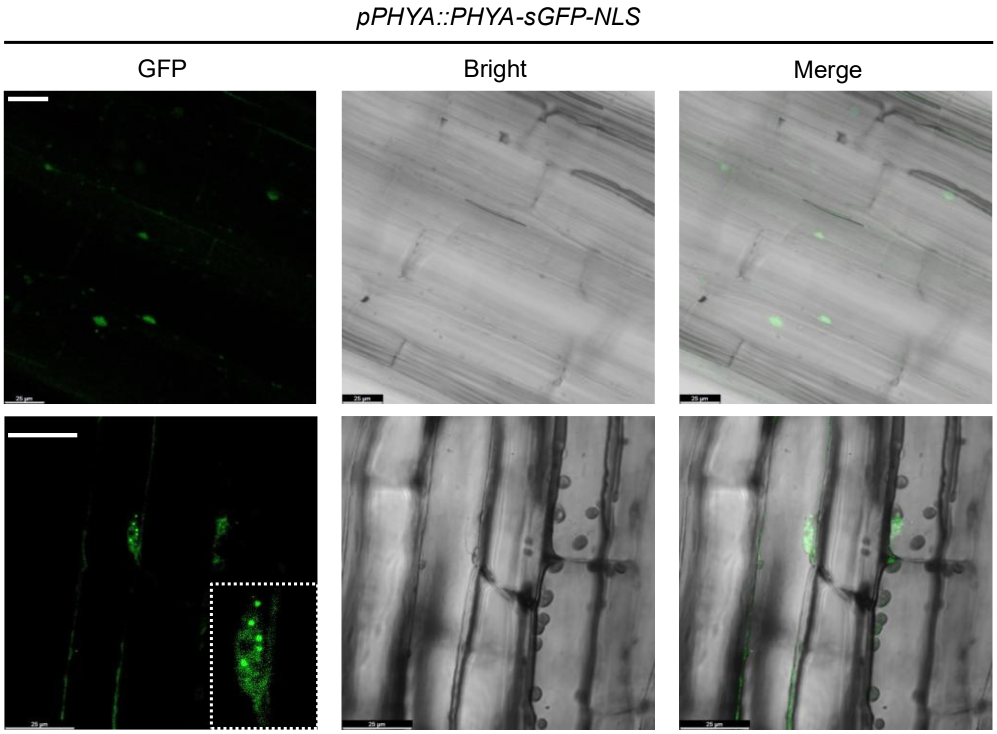


**Fig. S12. phyA-GFP-NLS accumulates in nuclei and photobodies in WLFR conditions.** Epidermal hypocotyl cell images from the transgenic line expressing *pPHYA::PHYA-sGFP-NLS*;*phyA-201*. GFP signals were detected using a 488 nm laser, and images were acquired at ZT7 on day six of the WLFR regime. The scale bar indicates 25 µm. Representative images from three independent replicates are shown.

**Fig. S13.**

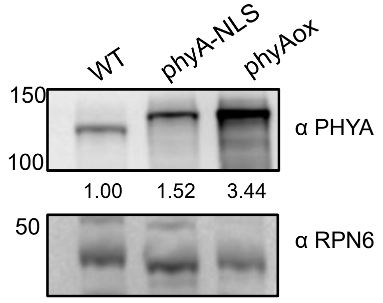


**Fig. S13. Comparison of phyA protein levels in the phyA-NLS and phyAox lines.** phyA levels in WT (Col-0), *pPHYA::phyA-sGFP-NLS* (phyA-NLS) and *p35S::phyA-RFP-terRbcS* (phyAox) determined by immunoblotting. Seedlings were grown in 8 h light with supplementary FR (PAR = 85 ± 5 μmol $m^{-2}$ $s^{-1}$, R:FR 0.20); 16 h dark photocycles (WLFR), at 22°C. Samples were collected at ZT7 on day 6. Anti-phyA antibodies were used for phyA detection, and an anti-RPN6 antibody was used as the internal control. The protein level quantification was normalised to WT in ImageJ. The WT recognized as 1.

**Table S1.**

| CONDATION | R/FR DEFINITION | | k1/k2 transition rate ratio according to oat phyA spectrum: | Pfr/Ptot according to oat phyA spectrum: | R:FR according to LI-COR spectrometer (SPC-00346) |
|---|---|---|---|---|---|
| | 640-700nm /700-760nm | 640-670nm /720-750nm | | | |
| WL | 0.9981 | 0.046775 | 3.7031 | 0.78737 | 8.5303 |
| WLFR | 1.8707 | 0.023234 | 0.37482 | 0.27263 | 0.2025 |

**Table S1.** R:FR calculations across commonly used waveband definitions, along with corresponding phytochrome transition rate and Pfr/Ptot predictions for conditions used in this work. The k1/k2 transition and Pfr/Ptot ratios were predicted according to Pr/Pfr conversion spectra of oat phyA (*55*). Calculations and scripts were obtained from Dr. Johanna Kramer.

### Table S2.

| Transgenic Materials | Primer Name | Sequence (5'- to -3') |
|---|---|---|
| *phyA-211* | phyA-F | GACACGATGATTCCTGCATC |
| | phyA-R | CAGCTGTGCGGTGCTCTAA |
| *phyA-201* | *phyA-201*-F | GAAGTGTTGACTGCTTCCACGAGT |
| | *phyA-201*-R | TAGCAAGATGCACAGAACGCC ( Need check with *Hrf1*) |
| PIF7ox | PIF7-F | ACTGCAAGTAAGGCGGATAAAGTC |
| | PIF7-R | GATCCATATTCATTGTCTGTGCG |
| *pif7-2* | pif7-2-F | GGAGAGCCATAGAGTTGG |
| | pif7-2-R | CGACATCTGAAACTGTTGC |
| | T-DNA _R | TAGCATCTGAATTTCATAACCAATCTCGATACAC |
| *pif7-1* | PIF7-1_F | CATCCTCTGGTTTATCCTATCACGCCG |
| | PIF7-1_R | CCGTTCATGGTCTAGGCG |
| | T-DNA_R | TGATAGTGACCTTAGGCGACTTTTGAACGC |
| NanoLUC | Nanoluc-F | ATGGTCTTCACACTCGAAGATTTC |
| | Nanoluc-R | TCAGTGATGGTGATGGTGATGGTG |
| NanoLUC -lgbit | Lgbit-F | TCTTCACACTCGAAGATTTCG |
| | Lgbit-R | GGAGGCTCGAGCGGTGCGATCGCCGCTTTGAGATATGCAGGTGT |
| NanoLUC-smbit | smbit-F | TCACCTGCATATCTCTAATGGTGACCGGCTACCGGCTGTTCG |
| | smbit-R | CGAGCGGTGCGATCGCCATGGAATAA |
| *pPIF7::PIF7-nanoLUC* | PIF7-Nluc-F | GGGGACAAGTTTGTACAAAAAAGCAGGCTTCAGTGATGCTCCATTTTGAAGAG TAC |
| | PIF7-Nluc-R | GGGGACCACTTTGTACAAGAAAGCTGGGTGATCTCTTTTCTCATGATTCGAAGA AC |

| Plasmid Constructs | Primer Name | Sequence (5'- to -3') |
|---|---|---|
| mUAV L0 | Aux-F | ATTACCGCCTTTGAGTGAGC |
| | Aux-R | TGCCACCTGACGTCTAAGAA |
| pMAP B L1 | PLX-F | GCGTAGAAACCAACATGC |
| | PLX-R | CAAAGGACCGCATGTCGCAAGTCG |
| *p35S::PIF7:sfGFP* | PIF7-F | TGGCTAGGTCTCCAATGTCGAATTAT |
| | PIF7-R | TGGCTAGGTCTCACGAACCATCTCTTTTCT |
| | sfGFP-F | GGTGGAGCAAGGGCGAGGAGCTGTT |
| | sfGFP-R | AAAGCTTACTTGTACAGCTCGTC |
| *p35S::PHYA:RFP* | RFP-F | ATGGTGTCTAAGGGCGAAGAG |
| | RFP-R | CTTGTACAGCTCGTCCATGCC |
| | phyA-F | GGTCTCTAATGTCAGGCTCTAGGCCGA |
| | phyA-R | GGTCTCACGAACTTGTTTGCTGCAG |
| Bifc:nYFP | n-YFP-F | ATGGTGAGCAAGGGCGAGGAG |
| | n-YFP-R | ATCCGCCACAACATCGAGTAG |
| Bifc:cYFP | c-YFP-F | GACAAGCAGAAGAACGGCATC |
| | c-YFP-R | TTACTTGTACAGCTCGTCCAT |
| *p35S::PHYA:RFP* | phyA-C | TCACCTGCATATCTCTAATG TTCAAGGATAGTGAA |
| | phyA-C | ACACCTGCATATCTCACGAAAGCTTCTAGTTCTTG |

| | | |
|---|---|---|
| | phyA-C1 | TCACCTGCATATCTCTAATGGAGAAAAAAATG |
| | phyA-C1 | ACACCTGCATATCTCACGAAAGCATTCCTC |
| | phyA-PAS | TCACCTGCATATCTCTAATGGTGACCAGTGAGATG |
| | phyA-PAS | ACACCTGCATATCTCACGAAACAGAAGACACCTGT |
| | phyA-HKRD | TCACCTGCATATCTCTAATGCGAACCGCAGTGAAG |
| | phyA-HKRD | ACACCTGCATATCTCACGAAGAAACTTGATTTCCC |
| *p35S::PIF7:nYFP* | PIF7-N | TCACCTGCATATCTCTAATGTCGAATTATGGA |
| | PIF7-N | ACACCTGCATATCTCACGAATCTAGCTGTCTTCAA |
| | PIF7-B | TCACCTGCATATCTCTAATGACCGGAGACAGAGAC |
| | PIF7-B | ACACCTGCATATCTCACGAATTGTGGCAAGTTGGC |
| | PIF7-C | TCACCTGCATATCTCTAATGACAAATGATGATTCCG |
| | PIF7-C | ACACCTGCATATCTCACGAAATCTCTTTTCTCATG |
| *p35S::*FHY1:EYFP/ *p35S::*FHY1:smbit | FHY1 | TCACCTGCATATCTCTAATGCCTGAAGTGGAAGTG |
| | FHY1 | ACACCTGCATATCTCACGAATTACAGCATTAGCGT |
| | EYFP | GTGAGCAAGGGCGAGGAGCTG |
| | EYFP | CTTGTACAGCTCGTCCATGCC |
| *p35S::*FHL:EYFP/ *p35S::*FHL:lgbit | FHL | TCACCTGCATATCTCTAATGGATGATGCAGATAAGA |
| | FHL | ACACCTGCATATCTCACGAATTACATCATGAGTGTAG |

**Table S2. Genotyping and Cloning primers used in this work.**

#### Table S3.

| Primer Name | Sequence (5’- to -3’) |
|---|---|
| PHYA-F | ATTGAAGGATGCTTGGATTTGG |
| PHYA-R | CCGTTACTCTTCATCATTACTTGAC |
| FHL-F | AAACCAGCGATGAAGCCTC |
| FHL-R | GCTTGATGTAGCAGCATAACC |
| FHY1-F (*35*) | GATGAAAGAGGAATCATCTGGA |
| FHY1-R: | AATCCTCTAAGTTCTGAGTCCCA |
| FHY3-F | TGGGAATCAACAAACAATGC |
| FHY3-R | AGAAATCTACTCCCTGTCCA |
| CHS-F | AGCTGATGGACCTGCAGGCATCTTGGC |
| CHS-R | TGCATGTGACGTTTCCGAATTGTCGAC |
| PP2AA3-F (*40*) | TAACGTGGCCAAAATGATGC |
| PP2AA3-R | GTTCTCCACAACCGCTTGGT |
| IAA29-F | CCGAATATGAAGATTGCGACAG |
| IAA29-R | GCAAAGATCTTCCATGTAACATCC |
| BIN2-F | GAGATGCCTGCTGCTGTAGT |
| BIN2-R | TGCTTGGAAAACGATCCCGA |
| CHS-(+)Gbox1-F (*32*) | CCCACCATTCAATCTTGGTAAG |
| CHS-(+)Gbox1-R | ACACCAACTTGGGTTTATTAGAG |
| CHS-(+)Gbox2-F (*32*) | TATTAGATTAGTAGGAGCTAATGATGGAGT |
| CHS-(+)Gbox2-R | TTATTATGTCTTAAGATACGTATCGCTTG |
| CHS-(-)Gbox-F | GTCTGCTCTGAGATCACAGCCG |
| CHS-(-) Gbox -R | GAGATGAGGCCGGGAACATCCT |
| FHY1-(+)Gbox -R (*32*) | GAGAGAGAGAGATAGAGAGAGTTCAA |
| FHY1-(+)Gbox -F | ACGCGCCAAATCAAACA |
| FHL-(+)Gbox -F | GGCCCACAATAGTCTCACTCC |
| FHL-(+)Gbox -R | ACCGTTGGCCCATGACAGA |

**Table S3. Real time q-PCR and ChIP-q-PCR primers used in this work.**
